# Engineering customized viral receptors for various coronaviruses

**DOI:** 10.1101/2024.03.03.583237

**Authors:** Peng Liu, Mei-Ling Huang, Hua Guo, Jun-Yu Si, Yuan-Mei Chen, Chun-Li Wang, Xiao Yu, Lu-Lu Shi, Qing Xiong, Cheng-Bao Ma, Fei Tong, Chen Liu, Jing Chen, Ming Guo, Jing Li, Zheng-Li Shi, Huan Yan

## Abstract

Coronaviruses display versatile receptor usage, yet in-depth characterization of coronaviruses lacking known receptor identities has been impeded by the absence of feasible infection models^1,2^. Here, we developed an innovative strategy to engineer functional customized viral receptors (CVRs). The modular design relies on building receptor frameworks comprising various function modules and generating specific epitope-targeting viral binding domains. We showed the key factors for CVRs to efficiently facilitate spike cleavage, membrane fusion, pseudovirus entry, and authentic virus propagation for various coronaviruses, resembling their native receptors. Applying this strategy, we delineated the accessible receptor binding epitopes for functional SARS-CoV-2 CVR design and elucidated the mechanism of entry supported by an amino-terminus domain (NTD) targeting S2L20-CVR. Furthermore, we created CVR-expressing cells for assessing antibodies and inhibitors against 12 representative coronaviruses from six subgenera, most of which lacking known receptors. Notably, a pan-sarbecovirus CVR supported entry of various sarbecoviruses, as well as propagation of a replicable HKU3 pseudovirus and the authentic strain RsHuB2019A^3^. Through combining an HKU5-specific CVR with reverse genetics, we successfully rescued and cultured wild-type and fluorescence protein-incorporated HKU5, a receptor-unidentified merbecovirus. Our study demonstrated the great potential of CVR strategy in establishing native receptor-independent infection models, paving the way for studying various viruses that are challenging to culture due to the lack of susceptible cells.

Graphic abstract

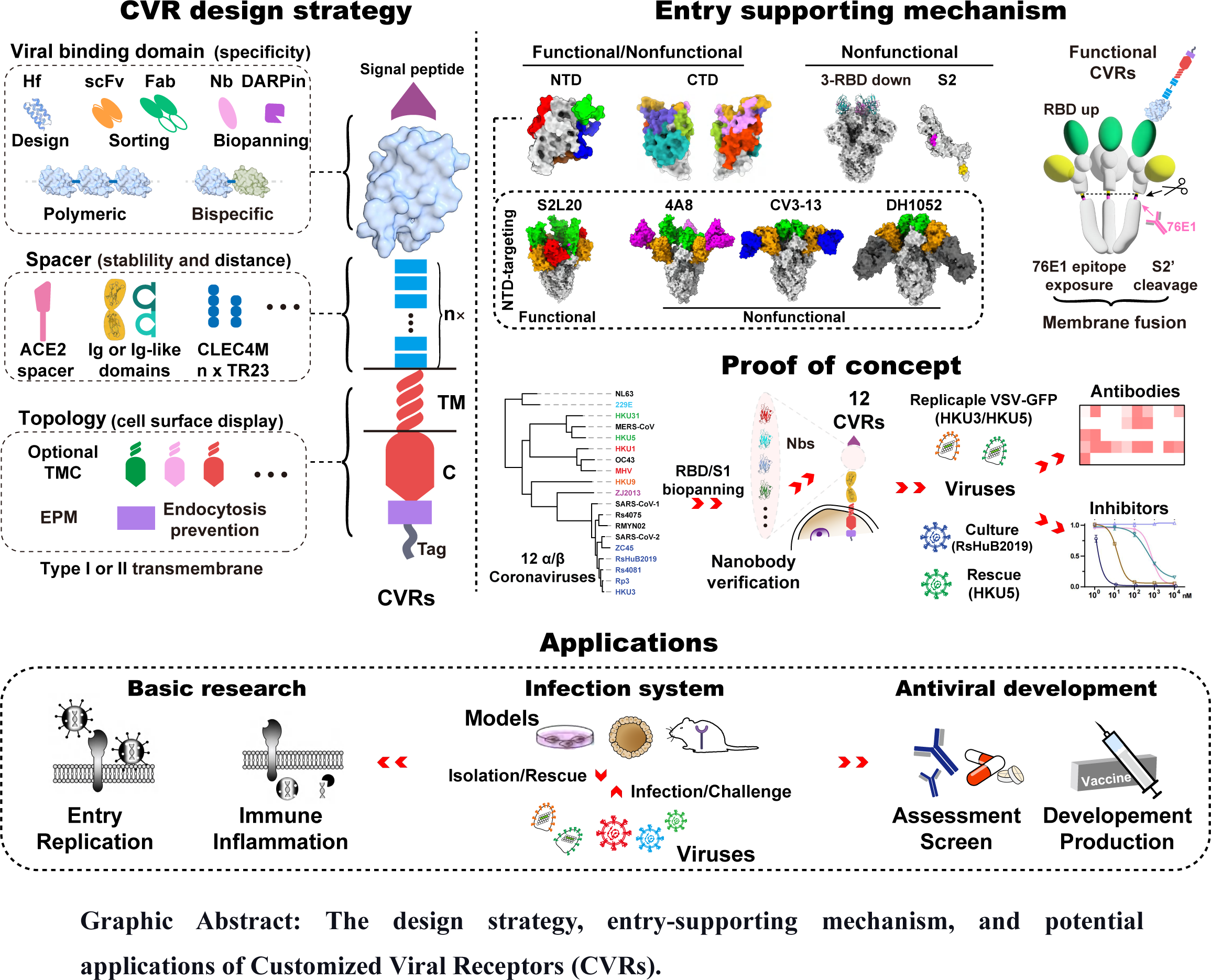

## Introduction

The *Coronaviridae* family encompasses hundreds of enveloped viruses categorized into four genera, α-, β-, γ- and δ-coronaviruses^4^. The emergence of human β-coronaviruses has led to three significant outbreaks in the 21^st^ century, highlighting the substantial zoonotic risks associated with various animal coronaviruses that are poorly studied, primarily infecting bats^5–8^.

Coronavirus entry is mediated by the trimerized spike (S) proteins. The full-length S may either remain intact or undergo cleavage by furin or other proteases at the S1/S2 cleavage site, yielding S1 and S2 subunits^9^. The S1 subunit engages in specific interaction with the receptor, leading to conformational changes that trigger membrane fusion mediated by the S2 subunit^10^. The activation of S2 fusion machinery is associated with the exposure and proteolytic of the S2’ cleavage site, which is right upstream of the fusion peptide (FP). A successful fusion involves a dramatic transition of the high-energy prefusion conformation to the low-energy post-fusion conformation of the spike trimer, with an extended intermediate that refolds and brings the two membranes into proximity to overcome the energy barrier for fusion^11^. Except for MHV, which naturally employs its amino-terminal domain (NTD) for receptor engagement, most coronaviruses use their carboxy-terminal domain (CTD) of the S1 subunit as their receptor binding domains (RBD), adopting either “down” or “up” conformations in the spike trimer^12^. The “up” conformation is believed to be more accessible for receptor engagement^13^. For example, SARS-CoV-2 recognizes the ACE2 protease domain (or head domain) through the extended receptor binding motif (RBM). The ACE2 binding mediated conformational change exposes the S2’ site, followed by proteolytic cleavage either by cell surface transmembrane serine protease 2 (TMPRSS2), the endosome-localized cathepsin L or other proteases^14^.

Coronaviruses can employ different receptors or adopt different receptor recognition mechanisms to utilize the same receptor^15–17^. Efforts in the past decades have led to the identification of four widely acknowledged protein entry receptors for coronaviruses: ACE2, Aminopeptidase N (APN), Dipeptidyl peptidase-4 DPP4, and mCEACAM1a^2^. TMPRSS2 has also been recently reported as an entry receptor for human coronavirus HKU1^18,19^. ACE2 represents the most extensively studied receptor supporting entry of various coronaviruses, including NL63, SARS-CoV-1, SARS-CoV-2, and several clades of bat sarbecoviruses and merbecoviruses^16,20–22^. Human ACE2 is an 805-aminoacid (aa) type I transmembrane protein consisting of signal peptide, head domain, neck domain, spacer sequences, transmembrane domain, and cytosolic domain. Cryo-EM structure demonstrated a dimerized structure, with a direct engagement of the head domain (protease domain) in SARS-CoV-2 RBD, particularly the α1 and α2-helix and the loop connecting the β3- and β4-sheets^23^. However, the contribution of other sequences for achieving the optimal receptor function remains not fully understood. Many alternative receptors capable of mediating SARS-CoV-2 entry have been reported, including CD147, AXL, KREMEN1, ASGR1, NRP1, CLEC4M, TMEM106B, etc^24–26^. However, their entry-supporting efficiency is generally low compared with the ACE2 receptor, probably due to the lack of evolutionary viral adaptation. Nevertheless, many coronaviruses do not use these reported receptors, and their receptor identity remains elusive. Numerous bat coronaviruses are known solely as sequences in databases, limiting our knowledge and countermeasures against these animal coronaviruses^1,5^.

Remarkably, many coronaviruses with unknown receptors often exhibited narrow cell tropism or a complete lack of known susceptible cells^1^. A primary challenge of conducting in-depth studies is the difficulty in culturing these viruses. Functional entry receptors are pivotal for establishing infection models for these viruses. However, the conventional strategy for native receptor identification is challenging and largely unpredictable. To address this unmet need, the alternative approach of establishing feasible infection models independent of native receptors is awaiting exploration, with few attempts on MHV reported for this purpose^27,28^.

The challenge of designing viral entry receptors with satisfied functionality is impeded by the lack of knowledge regarding the optimal viral surface to be targeted, and the critical sequence and structural requirements for achieving acceptable conformational changes coupling the downstream entry process. Notably, studies focusing on SARS-CoV-2 or MERS-CoV have elucidated several scenarios of ACE2 or DPP4-independent entry, either by alternative receptors, antibody-dependent FcγR-mediated entry, or membrane-anchored antibodies, either in a productive or nonproductive manner^24,25,29–32^. These findings indicate the specific ACE2-SARS-CoV-2 interaction is dispensable for viral entry, making it feasible to design receptors for various coronaviruses without known native receptor identities.

In this study, by dissecting the contributing sequences for ACE2 to support SARS-CoV-2 entry efficiently, we demonstrated that each part of the ACE2 contributes to the functionality in different ways. Nevertheless, all the ACE2 sequences were replaceable, enabling the rebuilding of various receptors with customized specificity by grafting viral binding domains generated by multiple methods. By deciphering the key factors affecting the functionality of the receptors, we developed a generally applicable modular design strategy to build functional customized viral receptors (CVRs) for supporting productive entry of viruses, either the vesicular stomatitis virus (VSV) based pseudoviruses or the authentic viral strains. Utilizing the engineered cell culture models expressing various CVRs, we demonstrated the advantage of this strategy in various applications, such as investigating viral entry mechanisms, assessing the efficacy and breadth of antibodies and other antivirals, improving coronavirus culture efficiency, and isolating or rescuing coronaviruses without known receptors.

## Results

### Modular design of customized viral receptors

We set out to delineate the role of each sequence or structural component of human ACE2 in functioning as an ideal receptor for SARS-CoV-2. We first tested the feasibility of using computationally designed ACE2-mimicking small helical frameworks to replace the ACE2 head domain while maintaining its receptor function. Four ACE2 chimeric proteins were created, with the head domain or head/neck domains replaced by two previously reported SARS-CoV-2 RBD binding helical frameworks, LCB1 and LCB3^33^. The SARS-CoV-2 authentic virus infection assays demonstrated that these chimeric proteins effectively supported viral infection (Extended Data Fig. 1).

We further investigated the importance of other ACE2 sequences by gradually reducing the remaining ACE2 components, including the neck domain, spacer, and cytosolic domain (D2-D5) (Fig. 1a). All chimeric proteins showed comparable SARS-CoV-2 RBD binding as examined by flow cytometry (Fig. 1b-c). However, the pseudovirus entry-supporting ability declined with decreasing ACE2 sequences, although the shortest 132aa protein maintained detectable receptor function, approximately 0.5% compared to the ACE2 group and 113-fold compared to vector control (Fig. 1d-e).

**Fig. 1.**
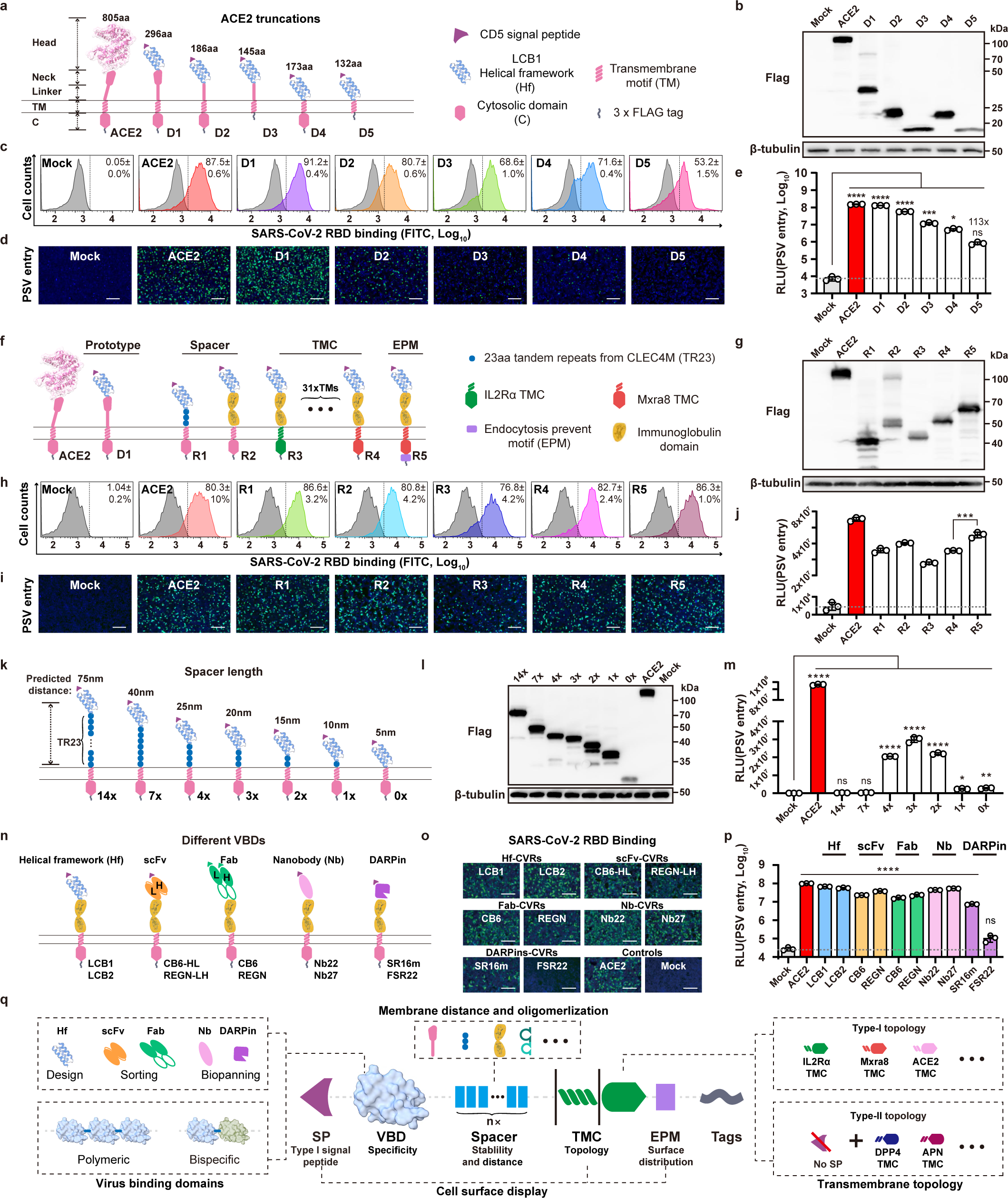
Modular design of customized viral receptors (CVRs) for efficient SARS-CoV-2 entry. **a-e,** Dissecting the importance of ACE2 sequences for its viral receptor function. **a,** Schematic representation illustrates the LCB1-ACE2 chimera with stepwise truncated ACE2 sequences. Protein expression levels (**b**) and SARS-CoV-2 RBD binding efficiency (**c**) in HEK293T transiently expressing the specified chimera. SARS-CoV-2 pseudovirus (PSV) entry in cells expressing the chimera was demonstrated by intracellular GFP (**d**) and RLU (**e**), respectively. **f-j,** Functionality of chimeric receptors with remaining ACE2 sequences substituted by domains from other proteins. **f,** Schematic representation delineates CVRs carrying exogenous spacer, transmembrane and cytosolic domain (TMC), and EPM sequences. The CVR expression (**g**), SARS-CoV-2 RBD-mFc binding (**h**), and PSV entry efficiencies (**i, j**) in HEK293T transiently expressing the indicated receptors. **k-m**, The impact of spacer length on CVR receptor function. Schematic representation illustrates CVRs with various TR23 tandem repeats, displaying predicted spacer length (**k**). CVR expression (**l**) and SARS-CoV-2 PSV entry efficiency (**m**) were evaluated in cells expressing the indicated CVRs. **n-p**, Different types of viral binding domains (VBDs) are compatible with CVR design. The SARS-CoV-2 RBD binding (**o**) and PSV entry (**p**) are supported by indicated CVRs transiently expressed in HEK293T cells. **q,** Schematic illustration of the modular design strategy for CVRs. RLU: relative light units. Scale bars: 200 μm. Data are presented as mean ± SD for n=3 biologically repeats for **c, e, h, j, m,** and **p.** One-way ANOVA analysis followed by Dunnett’s test for **e** and **m;** unpaired two-tailed Student’s t-tests for **j** and **p**.

Several chimeric proteins were subsequently designed with indicated domains replaced by corresponding sequences from other viral receptors or immune receptors (R1-R4), along with a construct carrying an endocytosis prevention motif (EPM) to enhance surface distribution (R5) (Fig. 1f)^34^. All chimeric proteins demonstrated well expression and efficient RBD binding (Fig. 1g-h). Particularly, the chimeric protein with the ACE2 neck domain substituted with triple (3×) 23aa tandem repeats (TR23) from CLEC4M or human IgG Fc supported efficient entry (Fig. 1i-j). Further substituting the remaining sequence with IL2Rα corresponding sequences maintained similar entry efficiency, suggesting that no ACE2-derived sequences are strictly required for SARS-CoV-2 entry (Fig. 1i-j). Among 31 tested transmembrane (TM) and several cytosolic domains from different receptors, the transmembrane and cytosolic domain (TMC) from the Chikungunya (CHIKV) receptor Matrix remodeling-associated protein 8 (Mxra8) exhibited the best performance (Extended Data Fig. 2)^35^. Constructs with EPM showed improved cell surface localization and enhanced entry-supporting ability (Fig. 1i-j, Extended Data Fig. 3). Additionally, constructs with a type-II transmembrane topology also efficiently supported SARS-CoV-2 and MERS-CoV entry, indicating the feasibility of both transmembrane topology for supporting coronavirus entry (Extended Data Fig. 4).

We then explored the impact of spacer length and oligomerization on entry-supporting efficiency by testing spacers with different copies of TR23 tandem repeats or immunoglobulin-like domains from human IgG or mCEACAM1a (Fig.1k-l, and Extended Data Fig. 5 and 6). Results indicated the triple TR23 or two immunoglobulin (Ig) or Ig-like domains represent the optimal spacer length for LCB1, while abolishing dimerization by Fc mutants has no significant impact on receptor function (Fig.1m and Extended Data Fig. 5 and 6) ^36^.

Subsequently, various SARS-CoV-2 RBD-targeting viral binding domains (VBDs) were tested for receptor grafting, including designed helical frameworks, designed ankyrin repeat proteins (DARPins), nanobody, scFv, and Fab (Fig. 1n and Extended Data Fig. 7). All these VBDs types are acceptable, with nanobodies showing superiority due to their small size, single-chain nature, and compatibility for bio-panning (Fig. 1o, p). We also demonstrated the functionality of a bi-specific receptor carrying two VBDs recognizing SARS-CoV-2 and MERS-CoV, respectively, and trimerized VBDs recognizing SARS-CoV-2 RBD (Extended Data Fig. 8). Additionally, we show the entry facilitated by soluble receptor adapters connecting viral RBD and ACE2 or FcγRIIa, respectively (Extended Data Fig. 9).

The functionality of CVRs compared with ACE2 was demonstrated through a series of experiments showing membrane fusion, authentic SARS-CoV-2 infection, and virus specificity in different cell types (Extended Data Fig.10). The entry-supporting efficiency of CVRs are significantly more efficient than several documented SARS-CoV-2 alternative receptors, coreceptors, entry factors, or binding proteins^24–26^ (Extended Data Fig.11).

Together, we proposed a modular design strategy for generating customized viral receptors to support efficient coronavirus entry, comparable in specificity and efficiency to their native receptors. A CVR prototype consisting of Type-1 transmembrane topology carrying signal peptide (SP), VBD, spacer, TMC, EPM, and C-terminal tags and its derivatives are delineated (Fig. 1q).

### Acceptable epitopes for functional CVRs

In our initial exploration of the relationship between CVR receptor function and binding affinity or neutralizing activity, we evaluated 25 neutralizing nanobodies targeting SARS-CoV-2 RBD. However, the results did not demonstrate a clear correlation between entry-supporting ability and binding affinity or neutralizing activity. These data underscores the influence of other critical factors, particularly the binding epitopes that are not clearly defined for the 25 nanobodies (Extended Data Fig.12).

Therefore, we engineered CVRs carrying scFvs derived from 22 well-characterized SARS-CoV-2 neutralizing antibodies (Abs) covering most reported neutralizing epitopes on NTD, CTD, or S2 (Fig. 2a)^37–44^. These antibodies were transformed into scFv-based VBDs with N-terminal heavy chain (HL) or N-terminal light chain (LH), resulting in 44 CVRs for evaluation (Fig. 2b). All CVRs were well-expressed and verified with SARS-CoV-2 spike trimer binding, except for 76E1 recognizing a hidden epitope exposed after receptor binding^42^ (Extended Data Fig.13). The scFv-CVRs recognizing epitopes close to the canonical RBM (sites i, ii and iii) supported efficient entry, while many other RBD core domain-targeting scFv also exhibited decent entry-supporting capabilities (Fig. 2c). However, not all RBD epitopes are suitable for CVR design, such as S309 and two antibodies recognizing a quaternary epitope spanning the dual-RBD interface that lock the spike in a closed conformation (BG10-19, S2M11)^37–39^. Unexpectedly, an S2L20-CVR recognizing an NTD epitope (site iv) showed potent entry-supporting ability, challenging the previous hypothesis that NTD neutralizing antibodies are insufficient to induce SARS-CoV-2 membrane fusion and entry in an ACE2-independent manner (Fig. 2c)^32,45^. We further demonstrated the expression, antigen binding, pseudovirus entry, and membrane fusion supported by ten selected CVRs (Fig. 2d).

**Fig. 2.**
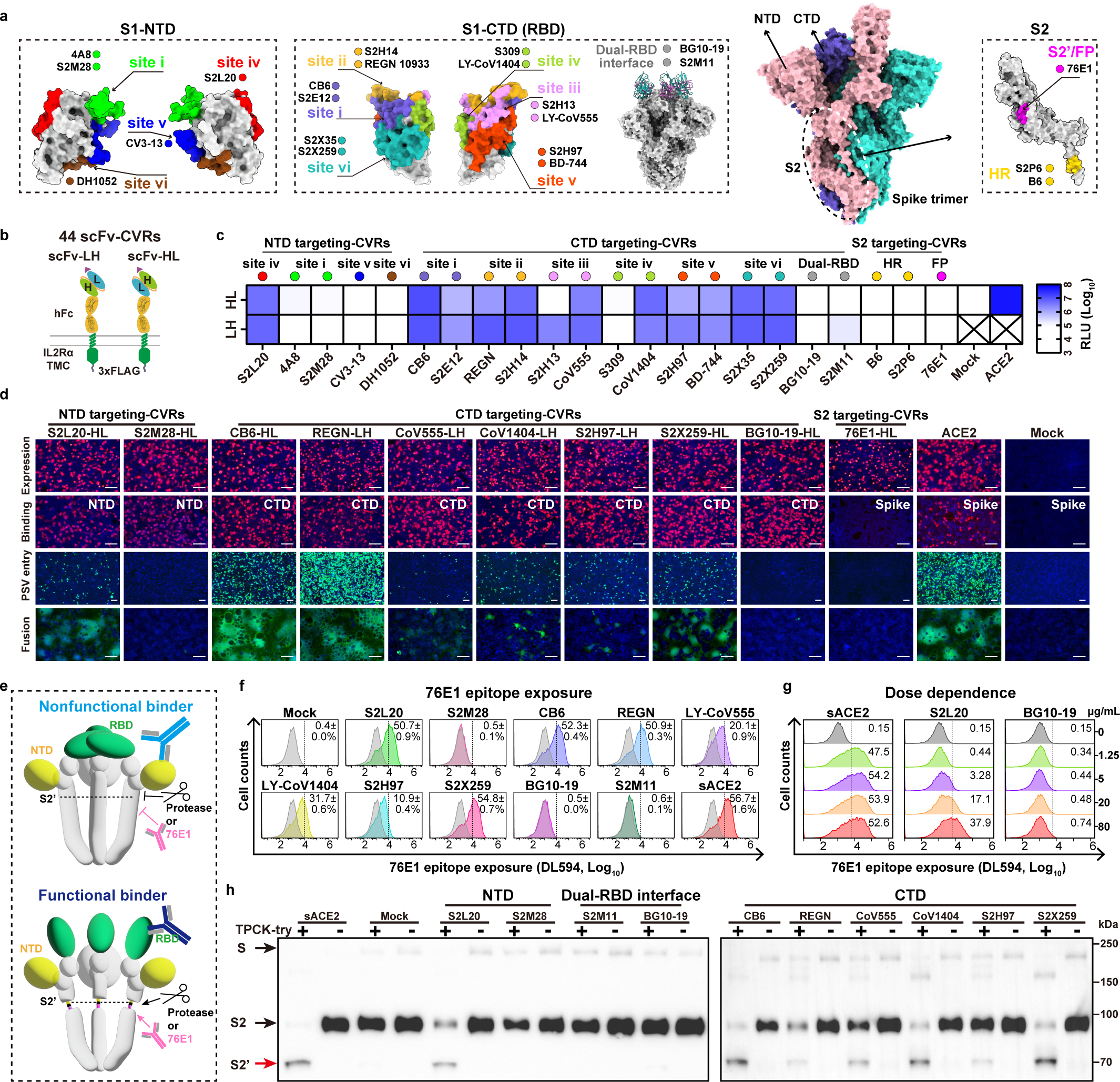
The impact of different binding epitopes on receptor functionality and the underlying mechanisms. **a,** Structural display of SARS-CoV-2 neutralizing epitopes in NTD, CTD, and S2 subunit, respectively. Various epitope types of 22 neutralizing antibodies are indicated. FP: fusion peptide. HR: heptad repeat. **b,** Schematic representation of 44 single-chain variable fragment (scFv)-CVRs with N-terminal light chain (LH) or heavy chain (HL), respectively. **c,** Heat map displaying SARS-CoV-2 PSV entry efficiency in HEK293T cells transiently expressing the indicated scFv-CVRs. **d,** Demonstration of CVR expression, antigen binding, PSV entry, and spike-mediated cell-cell fusion in HEK293T expressing representative scFv-CVRs. **e,** Cartoon elucidates the functional receptor-mediated RBD conformational change and the subsequent exposure of 76E1 binding epitope and cleavage at the S2’ site. **f,** Flow cytometry analysis of 76E1 epitope exposure in the presence of indicated soluble scFv-mFc recombinant proteins. Dashed lines denote thresholds for positive ratio calculation. sACE2: soluble ACE2 ectodomain. **g,** Dose-independent exposure of 76E1 epitope upon sACE2 or S2L20 coincubation, which was not detected in BG10-19. **h,** Trypsin-mediated cleavage of S2’ site in SARS-CoV-2 pseudovirus particles in the presence of scFv-CVRs with or without receptor function. TPCK-try: 10 μg/mL TPCK-treated trypsin. Scale bars: 100 μm. Data are presented as mean values for n=3 biologically repeats for **c**. Data are presented as mean ± SD for n=3 biologically repeats for **f**.

To elucidate why only specific epitopes are accessible for CVR design to realize receptor function, we proposed a hypothesis: CVR functionality is dependent on whether the interaction can induce a proper spike conformational change that leads to down-stream critical entry events required for membrane fusion, particularly the exposure and cleavage of S2’ cleavage site for fusion peptide activation^29,42,46^ (Fig. 2e). Consistently, although most of the tested scFv-mFc recombinant proteins can bind spike trimer, only scFv-mFc corresponding to the functional CVRs can induce the exposure of 76E1 epitope in a dose-dependent manner (Fig. 2e-g, and Extended Data Fig.14).

We further explored whether the exposure of the 76E1 epitope resulted in higher S2’ protease accessibility. After optimizing the experimental conditions for trypsin-based S2’ cleavage, we demonstrated that the ability of specific scFv-mFc to induce S2’ cleavage sensitivity aligns with the data from 76E1 epitope exposure assays (Fig. 2h and Extended Data Fig.15).

In summary, our data reveals that most CTD surfaces and specific NTD epitopes are accessible receptor binding motifs for generating functional CVRs, and the functionality is primarily determined by their capability to induce conformational changes capable of exposing the 76E1 epitope, which is subject to proteolytic cleavage at the S2’ site, thereby activating the fusion machinery.

### NTD-mediated sarbecovirus entry by S2L20-CVR

We next sought to characterize the NTD-mediated coronavirus entry facilitated by S2L20-CVR. We first confirmed that S2L20-CVR serves as a fully functional receptor for SARS-CoV-2, supporting membrane fusion, pseudovirus entry, and authentic virus infection (Fig. 3a). Additionally, S2L20-CVR effectively facilitated pseudovirus entry of the five SARS-CoV-2 variants of concern (VOCs) and the other three sarbecoviruses (BANAL-20-52, RaTG13, and GX-P2V) (Fig. 3b-d). As expected, SARS-CoV-1 and ZC45 cannot use S2L20-CVR for entry due to the lack of binding affinity (Fig. 3 c, d).

**Fig. 3.**
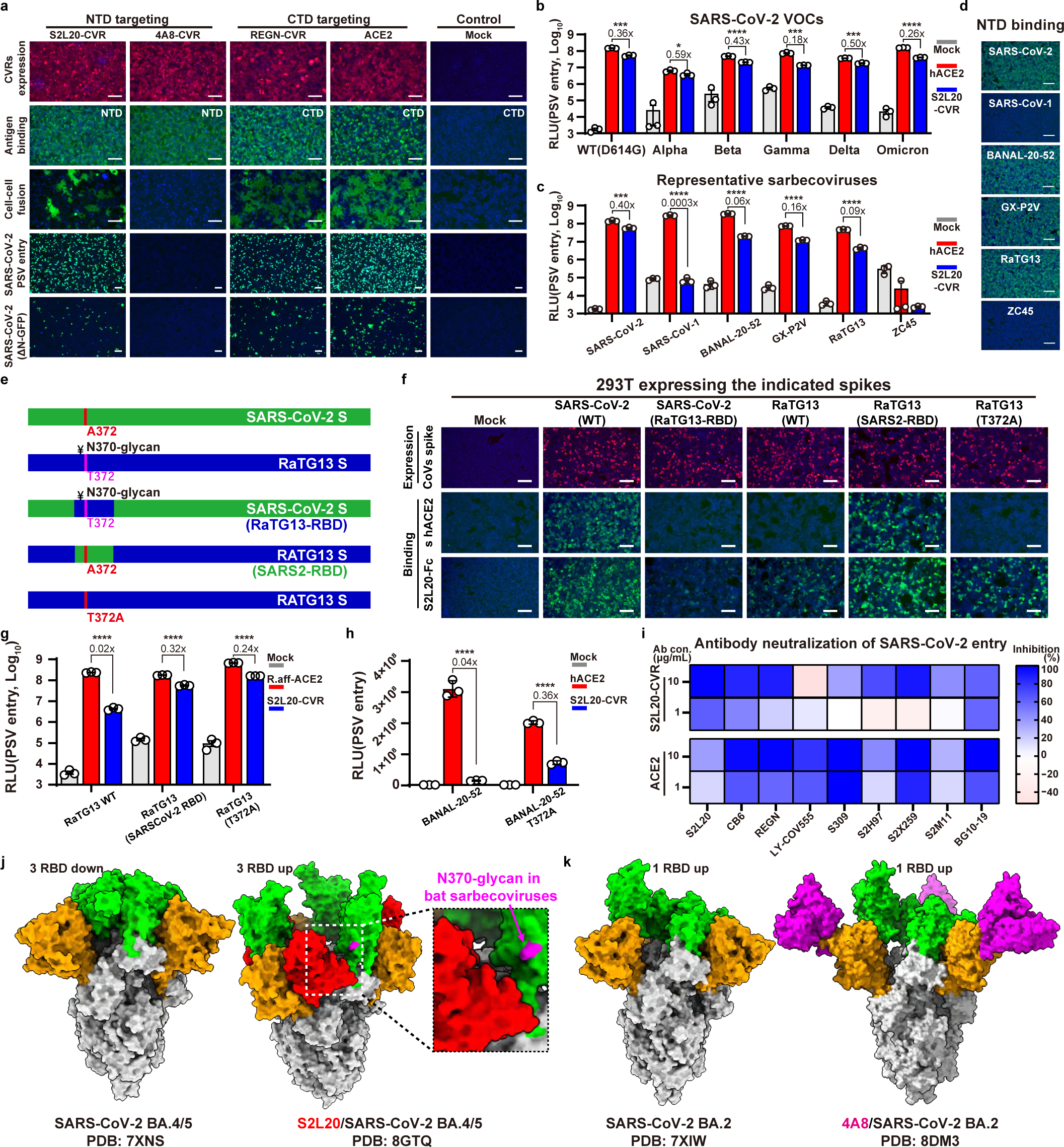
Characterization of NTD-mediated sarbecovirus entry supported by S2L20-CVR. **a,** SARS-CoV-2 binding, fusion, and viral entry supported by NTD-targeting S2L20-CVR. The CVR expression, NTD/CTD-mFc binding, cell-cell fusion, pseudovirus entry, and SARS-CoV-2(ΔN-GFP) infection in HEK293T stably expressing the indicated CVRs. **b,** PSV entry of SARS-CoV-2 VOCs in HEK293T expressing hACE2 or S2L20-CVR. **c, d,** The PSV entry (**c**) and NTD-mFc binding (**d**) efficiencies of various sarbecoviruses in HEK293T stably expressing hACE2 or S2L20-CVR. **e,** Illustration of the SARS-CoV-2 and RaTG13 RBD swap chimera. The residue usage in position 372, critical for the N370-glycosylation, is indicated. ¥: N-glycan. **f,** Spike proteins expression levels and the corresponding human ACE2 (shACE2) and S2L20-mFc binding efficiencies. **g, h,** Impact of T372A mutation on S2L20-CVR supported PSV entry of RaTG13 (**g**) and BANNAL-20-52 (**h**). **i,** Heatmap showing the inhibitory efficacy of indicated SARS-CoV-2 neutralizing antibodies against PSV entry in HEK293T-hACE2 or HEK293T-S2L20, with BSA as a control. **j,** Structures of SARS-CoV-2 BA.4/5 spike trimmer without antibody binding (left), or in complex with S2L20 (right). Dashed boxes highlighted the N370-glycan spatially proximate to the S2L20. **k,** Structures of SARS-CoV-2 BA.2 spike trimmers with (right) or without (left) the 4A8 binding. Orange: NTD; Green: CTD; Red: S2L20; Magenta: 4A8. The ratio of S2L20-supported entry compared to ACE2-supported entry is indicated in **c, g,** and **h**. Scale bars: 100 μm. Data are represented as mean ± SD with n=3 biological replicates for **b, c, g,** and **h**. Data representative of 2-3 independent experiments for **a-d, f-i**. Unpaired two-tailed Student’s t-tests for **c, b, g,** and **h**.

Despite showing similar NTD-binding efficiency, S2L20 showed much lower efficiency in supporting RaTG13 and BANAL-20-52 entry than SARS-CoV-2 (Fig. 3 c, d). Since the lack of an N370 glycan has been reported as a distinct feature of SARS-CoV-2, we generated CTD swap and point mutants to investigate the impact of CTD sequences and N370 glycan on S2L20-CVR dependent entry (Fig. 3e)^47^. Spikes carrying RaTG13 RBD or just a T372A mutation showed lower binding efficiency to the soluble forms of human ACE2(shACE2) or S2L20-mFc than those carrying SARS-CoV-2 RBD. Please note that RaTG13 has a lower affinity for hACE2 than its host’s ACE2 (*Rhinolophus. affinis* ACE2, R.aff ACE2) (Fig. 3f)^48^. The absence of the N370 glycan in SARS-CoV-2 due to a T372A mutation was hypothesized to interfere with S2L20 binding since this glycan is spatially close to the S2L20 after it binds to the NTD. Consistently, T372A mutation in the RaTG13 or BANAL-20-52 spike, abolishing the N370 glycosylation, significantly enhanced S2L20-CVR supported viral entry (Fig. 3g-h)^49^.

We next investigated whether SARS-CoV-2 CTD-targeting neutralizing antibodies could interfere with NTD-mediated entry in cells expressing S2L20-CVR compared to hACE2-expressing cells. As expected, S2L20 exhibited higher neutralizing activity in S2L20-CVR-expressing cells. Importantly, although several antibodies (LY-COV555, S309, and S2X259) showed reduced neutralizing efficiency in S2L20-CVR expressing cells, some CTD-binding antibodies exhibited similar neutralizing activity in both models (Fig. 3i).

These data suggest an association between RBD and S2L20-CVR mediated entry. Interestingly, the cryo-EM structure of S2L20 in complex with SARS-CoV-2 BA.5 revealed that S2L20 stabilizes the spike trimer in a three RBD “up” conformation, contrasting to the three RBD “down” conformation in BA.5 alone^50,51^. However, the binding of NTD-targeting antibodies, like 4A8, is unable to stabilize the RBD “up” conformation (Fig. 3j-k and Extended Data Fig. 17)^52^. We hypothesize the three RBD “up” conformation upon S2L20 binding may be crucial for S2L20-CVR receptor functionality. By contrast, mCEACAM1a, a receptor that binds to MHV spike trimer with three RBD down conformations, recognizes an NTD surface largely overlapped with the 4A8 epitope (Extended Data Fig. 17)^52–54^. This indicates different coronaviruses can adopt distinct receptor recognition mechanisms to achieve NTD-mediated entry.

### CVR supports efficient entry of various coronaviruses

The data presented above demonstrated the capability of CVRs to support efficient entry of ACE2 or DPP4-dependent coronaviruses. We extended our approach to generating CVRs capable of facilitating entry of 12 coronaviruses across the phylogeny, representing human, bat, and mouse coronaviruses from six distinct subgenera, the receptor for most of which remains unidentified^1^(Fig. 4a). To acquire acceptable VBDs for receptor grafting, we utilized magnetic beads and immunotube-assisted phage display biopanning to screen coronaviruses-specific nanobodies from the naïve libraries (Fig. 4b). Top CVR candidates, generated using a variety of nanobodies with validated RBD or S1 binding, demonstrated efficient pseudovirus entry for each coronavirus. Besides, we included a characterized broadly-neutralizing nanobody Nb27 for supporting RsHuB2019A. Binding kinetics of optimal nanobodies against the antigens from the 12 coronaviruses were determined through Bio-layer interferometry (BLI) assays (Fig. 4c and Extended Data Fig. 18). Efficient RBD or S1 binding and pseudovirus entry were demonstrated in 293T cells stabling expressing the indicated CVRs, achieving approximately 10^2^ to 10^4^-fold increase of entry compared with the mock control (Fig. 4d, e). Further examination of the CVR-supported entry of five different coronaviruses revealed that CVRs carrying EPM exhibited superior cell surface localization and higher entry-supporting ability (Extended Data Fig.19). Moreover, we verified the ability of several CVRs designed for 229E and MHV-A59 to support membrane fusion and authentic viral infection (Fig. 4f-h). Notably, both MHV-A59 NTD-targeting and CTD-targeting CVRs supported viral propagation, albeit with lower efficiency than the mCEACAM1a (Fig. 4i).

**Fig. 4.**
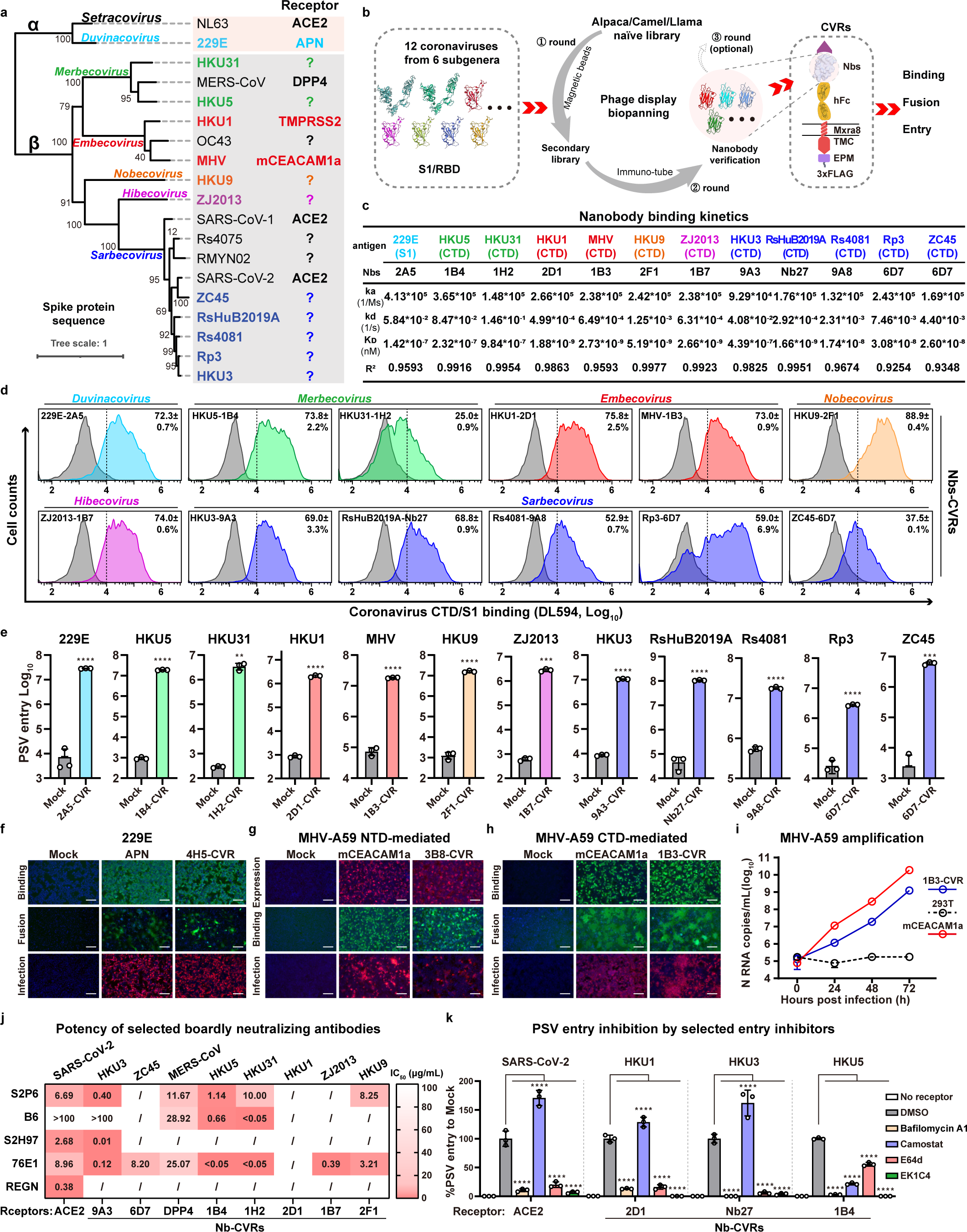
CVRs supported efficient entry of various coronaviruses with unidentified receptors. **a,** Phylogenetic tree based on spike protein amino acid sequences of representative coronaviruses. **b,** Workflow demonstrating the customization of nanobody-based CVRs for specific coronaviruses. **c,** Binding kinetics between optimal nanobodies and their respective antigens. **d,** Coronavirus CTD or S1 binding in HEK293T cells transiently expressing the corresponding CVRs. Dashed lines indicate thresholds for positive ratio calculation. **e,** PSV entry efficiencies of 12 representative coronaviruses in HEK293T cells transiently expressing the indicated CVRs. **f,** 229E S1-mFc binding, fusion, and authentic infection in HEK293T stably expressing APN or 4H5-CVR. **g-i,** Evaluation of NTD-mediated **(g)** or CTD-mediated **(h)** MHV antigen binding, cell-cell fusion, and authentic MHV infection in HEK293T cells stably expressing mCEACAM1a or the indicated CVRs. MHV-A59 RNA copies (N gene) in supernatant was quantified for infected cells expressing mCEACAM1a or 1B3-CVR (**i**). **j,** Summary of the IC_50_ of several broadly neutralizing antibodies against PSV entry of representative coronaviruses in HEK293T stably expressing the corresponding CVRs. The RBD-targeting REGN 10933 (REGN) was employed as a control. **/**: no inhibition detected. **k,** Inhibitory efficacy of inhibitors against PSV entry of SARS-CoV-2, HKU1, HKU3 and HKU5 in HEK293T cells stably expressing the indicated CVRs. Infection was examined by S protein immunofluorescence at 24 hpi for **f**, **g**, and **h**. Scale bars: 100 μm. Data are presented as mean ± SD and n=2-3 biologically repeats for **d, e, i,** and **k**. One-way ANOVA analysis followed by Dunnett’s test for **k;** unpaired two-tailed Student’s t-tests for **e**.

We next evaluated the CVR-based infection models for neutralizing antibody assessment, particularly anti-sera and broadly neutralizing antibodies. We compared the neutralizing activity of SARS-CoV-2 anti-sera, collected from COVID-19 convalescents or vaccinated individuals, on 293T cells expressing ACE2, LCB1-CVR recognizing the classical RBM, and Nb24-CVR recognizing an epitope distant from the RBM. The sera neutralization results based on the three receptors displayed a generally similar inhibitory profile, with Nb24-CVR showing slight differences (Extended Data Fig.20). This indicates the utility of the CVR-based system for evaluating the effectiveness of humoral immunity, ideally for CVRs recognizing the classical RBM region. We evaluated the cross-reactivity of several pan-β-CoV broadly neutralizing antibodies against several coronaviruses lacking conventional infection models, including antibodies recognizing an RBD site v epitope (S2H97), a stem region of the fusion machinery (S2P6, B6), and the S2/fusion peptide (76E1), with an RBM-targeting antibody REGN 10933 as a control. The results demonstrated that 76E1 exhibited the best breadth for cross-neutralization of the tested coronaviruses, consistent with sequence similarity of the recognized epitopes (Fig. 4j and Extended Data Fig.21).

Furthermore, we investigated the potential of the CVR-based infection system for evaluating other antivirals targeting different entry steps, including proteolytic cleavage, endosome acidification, and membrane fusion. Comparable inhibitory efficacy was observed when comparing infection models based on ACE2 or LCB1-CVR (Extended Data Fig.22). We further tested these inhibitors against SARS-CoV-2, HKU1, HKU3, and HKU5 entry in corresponding CVR-based infection models. Overall entry inhibitory efficiencies were similar among the four viruses, except for HKU5 displaying a higher sensitivity to TMPRSS2 inhibitor Camostat mesylate rather than the cathepsin inhibitor E64d (Fig. 4k). Our data confirmed the ability of CVR to support efficient entry for various coronaviruses. The novel infection models can be useful tools for assessing antibodies and other antiviral reagents against viruses lacking conventional infection models.

### Culture and rescue authentic coronaviruses through CVRs

To evaluate the capability of CVR-expressing cells to support multiple-round propagation of coronaviruses lacking known receptor identity, we utilized a reverse genetic system to generate propagation-competent VSV pseudoviruses with genomically encoded HKU3 or HKU5 spike proteins, replacing the VSV-G gene. A GFP-expressing cassette was additionally incorporated into the genome to facilitate visualization (Fig. 5a). Successfully rescued of the VSV-HKU3 and VSV-HKU5 was achieved with the aid of VSV-G proteins provided *in trans* (Fig. 5b). Following one round of amplification with VSV-G, infection was conducted in Caco2 cells in a VSV-G independent manner with or without the expression of Nb27, a pan-sarbecovirus CVRs recognizing the conserved site vi epitope on RBD (Fig. 5c-d and Extended Data Fig.23)^55^. Notably, efficient propagation of VSV-HKU3 and VSV-HKU5 can be observed in cells expressing the indicated CVRs, as evidenced by the syncytia formation and the accumulation of viral RNA in the supernatant, which was further enhanced by the exogenous trypsin treatment (Fig. 5d-g).

**Figure 5.**
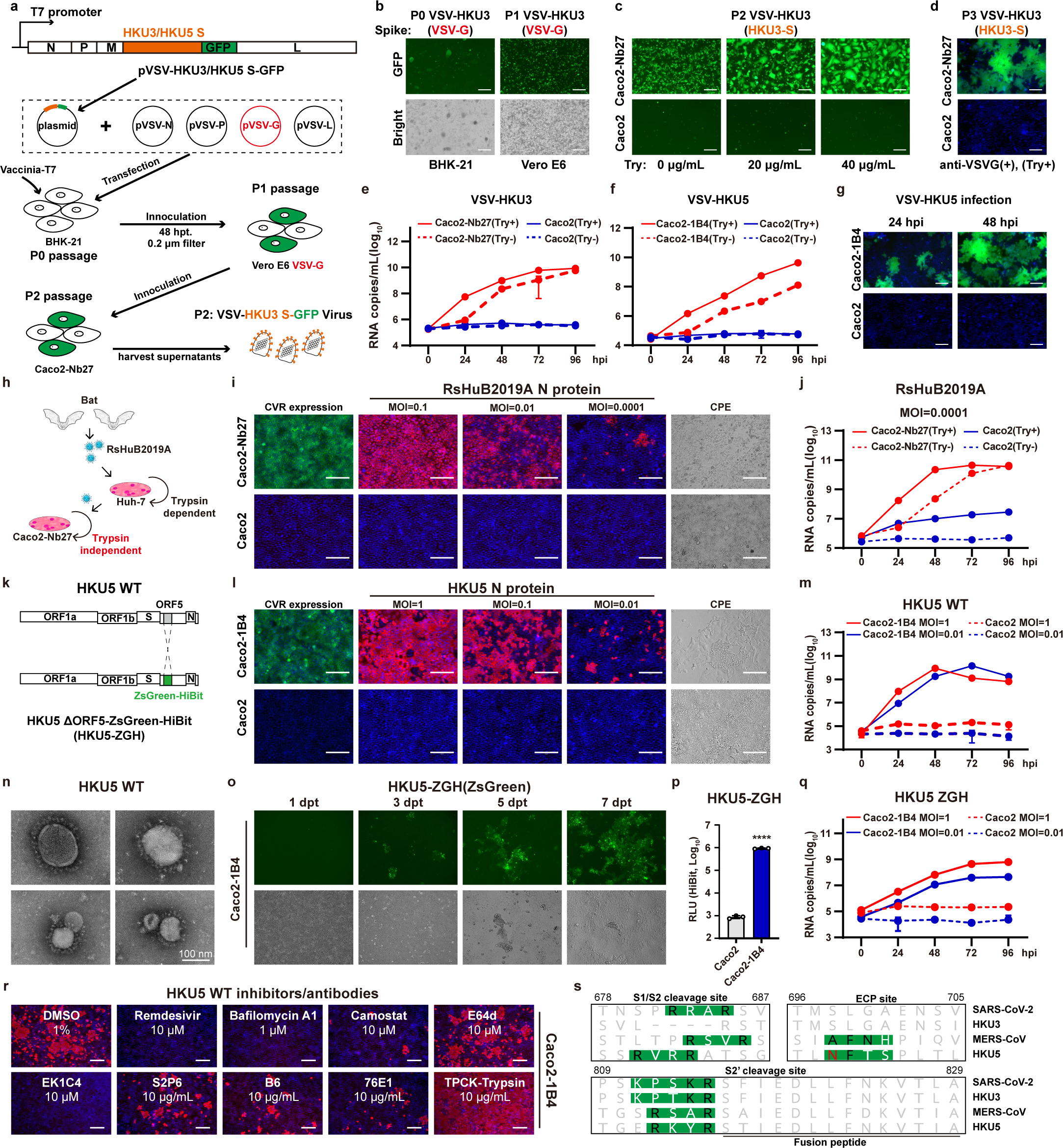
CVRs supported efficient propagation of VSV-based pseudotypes and authentic coronaviruses. **a,** Genetic organizations and workflow for generating replicable VSV-HKU3-GFP or VSV-HKU5-GFP. **b,** Successful rescue (P0) and amplification (P1) of VSV-HKU3-GFP assisted by VSV-G. **c, d,** Trypsin-enhanced cell-cell fusion (**c**) and VSV-G-independence (**d**) of VSV-HKU3-S-GFP infection in Caco2-Nb27 (MOI: 0.001). **e**, **f**, Accumulation of VSV-HKU3-GFP (**e**) or VSV-HKU5-GFP (**f**) RNA in the supernatant at indicated time points. **g,** VSV-HKU5-GFP mediated cell-cell fusion in Caco2 or Caco2-1B4 cells at MOI=0.1. **h,** Cartoon illustrating trypsin-independent propagation of RsHuB2019A in Caco2-Nb27 cells as compared with trypsin-dependence in Huh-7. **i**, CVR expression (green), RsHuB2019A N protein (red), and cytopathic effect (CPE, bright field) in cells inoculated with RsHuB2019A at indicated MOI (no trypsin). **j**, Accumulation of viral RNA in supernatant of cells infected with RsHuB2019A with or without trypsin. **k,** Genetic organizations of the HKU5 ΔORF5-ZsGreen-HiBit (HKU5-ZGH). **l,** CVR expression (green), N protein (red), and the CPE in indicated cells inoculated with HKU5 at different MOI. **m,** Accumulation of HKU5 RNA in supernatant of cells inoculated with HKU5 at different MOI. **n,** Transmission electron microscopy analysis of HKU5-WT virions. **o-q,** Increase in ZsGreen intensity (P0) (**o**), ZsGreen-HitBit signal (**p**), and supernatant RNA copies of HKU5-ZGH (**q**) in Caco2-1B4 cells. **r,** Efficacy of indicated antiviral reagents against HKU5 infection in Caco2-1B4 cells assessed by intracellular N proteins at 48 hpi. **s,** Overview of the protease cleavage sites of selected coronaviruses. The residue responsible for reduced endosomal cysteine protease activity (ECP) is marked in red, numbering based on SARS-CoV-2. Scale bars: 125 μm for **i, l,** and **o,** and 100 μm for **b-d, g,** and **r**. Data are presented as mean ± SD for n=2 biologically repeats for **e, f, j, m,** and **q**. Unpaired two-tailed Student’s t-tests for **p**.

Subsequently, we investigated whether CVRs can facilitate efficient propagation of authentic coronavirus requiring strict culture conditions. RsHuB2019A, a relative of HKU3, is an ACE2-independent bat sarbecoviruses recently isolated from field samples^3^. Isolation and propagation of this virus was carried out in Huh-7 under a serum-free culture condition with exogenous trypsin, with viral infection being difficult to detect while maintaining normal cell morphology (Fig. 5h). Our results demonstrated that the Caco2 cells stably expressing Nb27-CVR (Caco2-Nb27) efficiently supported RsHuB2019A propagation, even at very low MOIs (Fig. 5i and Extended Data Fig.24). Unlike Huh-7, Caco2-Nb27 supported efficient RsHuB2019A propagation in trypsin-free culture medium supplemented with 2% FBS, enabling observation of cytopathic effect (CPE) (Fig. 5i-j and Extended Data Fig.25).

Furthermore, we explored the feasibility of rescuing a representative bat merbecovirus, HKU5, by combining reverse genetics and the novel CVR-based infection system. Thus, we generated a full-length infectious clone of wild-type (WT) HKU5, along with a fluorescence protein with ORF5 substituted by a ZsGreen-HiBit reporter (ZGH) (Fig. 5k). Utilizing the same cell line for VSV-HKU5 propagation (Caco2-1B4), we successfully rescued both the WT and the ZGH version of HKU5 authentic viruses. Efficient amplification was observed in cells inoculated with HKU5 at different MOIs, as indicated by the nucleocapsid (N) protein immunostaining and the accumulation of genomic RNA in the supernatant over time (Fig. 5l-m). Electron microscopy revealed typical morphology of “crown-shaped” virions with diameters of approximately 100 nm (Fig. 5n). Although the HKU5-ZGH exhibited relatively slow amplification kinetics, likely due to the deleted ORF5 and foreign gene insertion, the expression of ZGH facilitated real-time visualization and quantification of viral amplification (Fig. 5o-q). Consistent with previous reports, HKU5 can amplify in Vero-E6 cells only in the presence of exogenous trypsin (Extended Data Fig.27)^56^.

Lastly, we assessed several antiviral reagents against HKU5 infection in Caco2-1B4 cells. Immunostaining of N protein revealed that trypsin significantly enhanced the infection, while most inhibitors blocked HKU5 infection. Consistent with the pseudovirus entry assay data (Fig. 4j), HKU5 infection was inhibited by Camostat mesylate but not E64d, further demonstrating the TMPRSS2 dependence for HKU5 infection (Fig. 5r and Extended Data Fig.26). This protease preference is in line with the sequence features at the critical cleavage sites for HKU5 (Fig. 5s)^56^.

## Discussion

The elusive identity of the receptor used by many coronaviruses presents substantial challenges in comprehending their life cycle and spillover risk, particularly those phylogenetically related to known high-risk β-coronaviruses. Lessons learned from the COVID-19 pandemic underscore the urgent imperative to study these viruses to prepare for future outbreaks. However, in-depth research and vaccine/antiviral development for these viruses are hindered by the lack of feasible infection models for virus isolation and culture^1^. Here, we proposed a novel strategy for customizing functional viral receptors for various coronaviruses, especially those lacking known native receptor identities. Our approach involves the modular design of CVRs in a single open-reading frame format, enabling molecular grafting of viral binding domains (VBDs) customized for specific viral surface epitopes. By targeting conserved epitopes compatible with CVR design, CVRs could potentially exhibit a broad recognition spectrum for coronaviruses from distinct clades or lineages.

We demonstrated that the VBDs can adopt various structures with binding affinity to viral surface proteins. Compared to Fab fragments, single polypeptide chain structures like scFv or nanobody are more suitable modules for CVR design and are compatible with the library biopanning system. The excellent performance of helical framework or DARPin-based CVRs, highlights the potential of computational de novo design of VBDs for various viruses. Besides efficient binding, maintaining an optimal distance between VBD and the cell membrane is critical for CVR functionality, although this distance may vary for spacers with distinct structures, orientations, and flexibility. Additionally, we demonstrated the soluble adaptor strategy in supporting viral entry, realized by a bio-specific adaptor retargeting the viruses to a cell surface-expressed receptor, such as ACE2.

Additionally, the functionality of the CVR highly depends on the acceptable epitopes recognized by the VBDs. The ability of the specific viral surface regions to serve as functional receptor binding motifs likely depends on whether a VBD recognizing this region can induce proper conformational changes leading to membrane fusion. Therefore, CVRs targeting S2, most epitopes of NTD, and some epitopes on CTD are nonfunctional. We revealed a close link between the CVR functionality and their ability to induce the exposure of 76E1 epitope, encompassing the critical S2’ cleavage site and part of the fusion peptide. However, the conformational change crucial for exposing this epitope remains unclear, although a transition of the RBD from the “down” to “up” conformation seems crucial^13^. Consistently, CVR using antibodies recognizing the three RBD “down” epitopes and locking the spike in this conformation showed no entry-supporting ability^38,39^. Future structural studies could be conducted to elucidate this critical event.

There appears to be no restriction for coronaviruses to employ their NTD or RBD for receptor engagement, as exemplified by the receptor function of NTD-targeting S2L20-CVR. Additionally, we also showed that MHV infection can be efficiently supported by either NTD or CTD-targeting CVRs, suggesting the possibility of MHV, or its relatives, recognizing an alternative receptor through their CTD^57^. It is also possible that SARS-CoV-2, or other sarbecoviruses, may develop NTD-mediated entry in the previous or future evolution. Notably, an infection-enhancing antibody targeting NTD of SARS-CoV-2, DH1052^58^, was unable to be utilized to build a functional CVR in our study. These indicate differences in mechanisms between the soluble antibody-mediated antibody-dependent enhancement (ADE) and membrane-anchored CVRs-mediated viral entry.

Our CVR strategy allows the modular design of customized receptors to manipulate cell susceptibility to specific viruses. This approach enables the isolation or rescue of coronaviruses regardless of receptor identity or conventional susceptible cells. Overcoming limitations of native receptors, such as enhancing affinity, altering epitopes, adjusting specificity, changing structures, or getting rid of physiological function interference, underscores the potential of this strategy. However, several limitations should be noted when employing CVR-based models. Differences in targeted epitopes and protein structures may result in inconsistencies when assessing RBM-targeting neutralizing antibodies or sera. Additionally, slight differences in the entry pathway may exist for some CVRs that trigger a conformational change different from those induced by native receptors. It is noteworthy that CVR transgenic mice might be useful for evaluating viral pathogenesis and vaccine/antiviral protection *in vivo.* However, the variations in tissue expression patterns may limit the mimicking of natural infection.

To our knowledge, this study demonstrated the first case of rescuing and culturing a coronavirus without known receptor identity based on a genetically modified cell culture model independent of native receptors. Our findings pave the way for the rapid design of novel viral infection models for difficult-to-culture viruses, including those beyond coronaviruses, facilitating further advances in basic research, antiviral therapeutics, and vaccine development.

## Author contributions

H.Y., P.L., M.L.H., H.G., and J.L. conceived the study and designed the experiments; P.L., M.L.H., H.G., J.Y.S., Y.M.C., C.L.W., X.Y., C.L., L.L.S. and J.L. established the assays and methods, designed and cloned receptor and protein constructs and conducted related experiments; P.L., M.L.H., L.L.S., C.B.M., Q.X., F.T. and C. L. screened the nanobodies; P.L., M.L.H., L.L.S. conducted 229E and MHV authentic viruses related experiments at Wuhan University; P.L. and H.G. conducted the RsHuB2019A and HKU5 authentic viruses and HKU5 negative staining for EM imaging in Wuhan Institute of Virology; J.C. and P.L. constructed the infectious clones of HKU5 and HKU5-ZGH, respectively. M.G. conducted the SARS-CoV-2 authentic viruses infection in ABSL-III in Wuhan University; H.Y., P.L., M.L.H., C.L.W., and X.Y. analyzed the data; H.Y., and P.L., M.L.H. prepared the original draft of the manuscript, H.Y., Z.L.S., P.L. revised the manuscript with input from all authors. H. Y. and Z.L.S. funded this study. H. Y. supervised the project.

## Acknowledgments

We are grateful for the funding support from the National Key R&D Program of China (2023YFC2605500 to H.Y. and Z.L.S.). National Natural Science Foundation of China (NSFC) projects (82322041, 32270164, 32070160 to H.Y., 32300141 to H.G.), Fundamental Research Funds for the Central Universities (2042023kf0191, 2042022kf1188 to H.Y.), and Natural Science Foundation of Hubei Province (2023AFA015 to H.Y.), and China Postdoctoral Science Foundation (2023M733708 to H.G.). We gratefully acknowledge Xiang-Xi Wang (Institute of Biophysics, Chinese Academy of Sciences) for providing the human sera collected post-vaccination (SARS-CoV-2 CoronaVac, Sinovac) and collected by Beijing Youan Hospital (approval number LL-2021-042-K), and Kun Cai (Hubei Provincial Center for Disease Control and Prevention) for providing sera collected from Wuhan COVID-19 convalescents (identification number 2021-012-01).

We gratefully acknowledge Zhao-Hui Qian (Chinese Academy of Medical Sciences and Peking Union Medical College) for providing BANNAL-20-52 related spike expressing plasmids. We also thank Pei Zhang and Bi-Chao Xu (Wuhan Institute of Virology) for their help with the ultracentrifugation and Electron Microscopic analysis. We thank Ying-Jian Li (Wuhan University) for assisting in establishing the yeast cloning methods. We also want to express our gratitude to the ABSL-3 facility and other core facilities of the Key Laboratory of Virology, Wuhan University.

## Data and code availability

This study did not generate custom computer code.

Any additional information required to reanalyze the data reported in this paper is available from the lead contact upon request.

## RESOURCE AVAILABILITY

### Lead contact

Further information and requests for resources and reagents should be directed to and will be fulfilled by the lead contact, Huan Yan

### Materials availability

All reagents generated in this study are available from the lead contact with a completed Materials Transfer Agreement.

## Methods

### Cell lines

HEK293T (CRL-3216), Vero E6 (CRL-1586), A549 (CCL-185), BHK-21 (CCL-10), Caco-2 (HTB-37), Neuro2a (CCL-131) and the bat epithelial cell line Tb 1 Lu (CCL-88) were purchased from the American Type Culture Collection (ATCC). The human hepatocellular carcinoma cell line Huh-7 (SCSP-526) was obtained from the Cell Bank of Type Culture Collection, Chinese Academy of Sciences. All cells were maintained in Dulbecco’s Modified Eagle Medium (DMEM, Monad), supplemented with 10% fetal bovine serum (FBS). Additionally, an I1-hybridoma cell line (CRL-2700), producing a neutralizing mouse monoclonal antibody against VSV-G, was cultured in minimum essential medium with Earle’s balanced salts solution, 2 mM L-glutamine (Gibco), and 5% FBS. All cell lines were incubated at 37℃ in 5% CO_2_ with regular passage every 2-3 days.

### Virus and host gene sequences

All viral genome or gene sequences were sourced from GenBank or GISAID databases with the following accession numbers. Viruses: SARS-CoV-1 (NC_004718), SARS-CoV-2 (NC_045512), MERS-CoV (NC_019843), HKU3 (DQ022305), Rp3 (DQ071615), HKU5 (NC_009020), HKU31 (MK907286), HKU9 (NC_009021), Zhejiang2013 (NC_025217), Rs4081 (KY417143), MHV-A59 (NC_048217), NL63 (JX504050), 229E (OQ920101), HKU1 (NC_006577), OC43 (AY391777), RmYN02 (EPI_ISL_412977), ZC45 (MG772933), RsHuB2019A (OQ503498). The spike protein for Rs4075 (KC880993). Receptors: ACE2 (NM_001371415), R.affinis ACE2 (MT394208), DPP4 (NM_001935), APN (NM_001150), mCEACAM1a (NM_001039186), AXL (NM_001699), NRP1 (NM_001024628), SCARB1 (BC143319), KREMEN1 (NM_032045), ASGR1 (NM_001671), CD147 (AB085790), CLEC4M (KJ902090), LRRC15 (NM_001135057)^59^, TMEM106B (NM_018374) ^60^, TMPRSS2 (NM_001135099). All receptor and viral gene sequences utilized in this study were commercially synthesized by Genewiz or GenScript.

### Plasmids

All plasmids expressing type-I transmembrane CVRs were constructed by inserting the human codon-optimized CVR sequences into a lentiviral transfer vector (pLVX-EF1a-Puro, Genewiz) with an N-terminal CD5 secretion leading sequence (MPMGSLQPLATLYLLGMLVASVL) and C-terminal 3×FLAG tag (DYKDHD-G-DYKDHD-IDYKDDDDK). For the type-II transmembrane CVRs, the C-terminal ectodomains were replaced by corresponding CVR modules, along with a C-terminal 3×FLAG tag. Chimeric protein-coding sequences were generated using overlapping PCR, direct sequence synthesis, or restriction endonuclease digestion and ligation.

Plasmids expressing the Spike protein of various coronaviruses for VSV pseudotyping were constructed by inserting human codon-optimized spike coding sequences into either the pCAGGS vector or pcDNA3.1(-) vectors with C-terminal 13-18 residues substituted with an HA tag (YPYDVPDYA) to enhance VSV pseudotyping efficiency and facilitate detection^61^. Several spike genes were also introduced into the pLVX-IRES-ZsGreen vectors for flow cytometry-related assays, including the scFv-mFc binding and the 76E1 epitope exposure assays.

Plasmids expressing secreted fusion proteins, such as coronavirus antigen-Fc, scFv-Fc, and nanobody-Fc, were constructed by inserting the coding sequences into pCAGGS. These constructs featured an N-terminal CD5 secretion leading sequence (MPMGSLQPLATLY LLGMLVASVL) and a C-terminal Twin-Strep Tag II following 3×FLAG tandem sequences (WSHPQFEKGGGSGGGSGGSAWSHPQFEK-GGGRS-DYKDHDGDYKDHDIDYKDDDDK) for purification or detection. Plasmids encoding codon-optimized anti-ACE2 antibodies H11B11^62^, B6, S2P6, 76E1, S2H97, and REGN10933 were constructed by integrating the heavy-chain and light-chain coding sequences into pCAGGS with an N-terminal CD5 leader sequences. For DSP-based cell-cell fusion assays, the split protein genes were inserted into pLVX-EF1a-Puro. The coding sequences for the dual reporter split proteins, namely RLuc (1-155)-sfGFP (1-157) and sfGFP (158-231)-RLuc (156-311), are previously descirbed^16^.

### Stable cell lines

Cells stably expressing distinct CVRs and other receptors were established through lentivirus transduction and subsequent antibiotic selection. Lentiviruses carrying the target genes were generated by co-transfecting lentiviral transfer plasmid (pLVX-EF1a-Puro) with packaging plasmids pMD2G (Addgene, 12259) and psPAX2 (Addgene, 12260) into HEK293T cells through GeneTwin transfection reagent (Biomed, TG101). The lentivirus-containing supernatant was harvested and pooled at 24 and 48 hours post-transfection. Cell transduction was carried out in the presence of 8 μg/mL polybrene. Stable cell lines were selected and maintained in a growth medium supplemented with puromycin (1 μg/mL). Generally, cells exhibiting stability for at least ten days were utilized in subsequent experiments.

### SARS-CoV-2 reactive antisera

SARS-CoV-2 antisera were obtained from vaccinated individuals (SARS-CoV-2 CoronaVac, Sinovac), approximately 21 days post-vaccination and Wuhan COVID-19 convalescents around one year post-infection, respectively. Ethical approval for the vaccinated individuals was granted by the Ethics Committee (seal) of Beijing Youan Hospital, Capital Medical University, with approval number LL-2021-042-K. The collection of sera from Wuhan COVID-19 convalescents was conducted in collaboration with the Hubei Provincial Center for Disease Control and Prevention and Hubei Provincial Academy of Preventive Medicine (HBCDC), following written consent and under the approval of the Institutional Review Boards with the identification number 2021-012-01. Sera were heat-inactivated at 56℃ for 30 minutes.

### Bioinformatic and computational analyses

Multi-sequence alignment was analyzed by Geneious Prime software or mafft (v7.407) with default parameters. Phylogenetic trees were constructed by IQ-TREE (http://igtree.cibiv.univie.ac.at/) with the WAG substitution model (1000 Bootstraps) and polished with iTOL (v6) (http://itol.embl.de). The cryo-EM structures were displayed and marked by ChimeraX with PDB accession numbers indicated in figures or legends.

### Protein expression and purification

The proteins for binding, neutralizing, or biopanning-related assays were produced in HEK293T by transient transfection with plasmids using GeneTwin reagent (Biomed, TG101-01), following the manufacturer’s guidelines. Protein-containing supernatants were harvested every 2-3 days post-transfection, pooled, clarified, and proceeded to purification. Proteins fused with Fc were captured using Pierce Protein A/G Plus Agarose (Thermo Scientific, 20424), eluted with pH 3.0 glycine (100 mM in H_2_O), and immediately pH-balanced by 1/10 volume of UltraPure 1 M Tris-HCI, pH 8.0 (15568025, Thermo Fisher Scientific). Proteins with Twin-Strep Tag II were enriched using Strep-Tactin XT 4Flow high-capacity resin (IBA, 2-5030-002), washed, and eluted with buffer BXT (100 mM Tris/HCl, pH 8.0, 150 mM NaCl, 1 mM EDTA, 50 mM biotin). The eluted proteins were concentrated and buffer-exchanged to PBS through ultrafiltration, aliquoted, and stored at −80℃. Protein concentrations were determined using the Omni-Easy Instant BCA Protein Assay Kit (Epizyme, ZJ102).

### Western blot

For detecting the cellular expression of CVRs or other receptors, cells were washed once with PBS and lysed using RIPA buffer (50 mM Tris-pH 7.4, 150 mM NaCl, 1%TritonX-100, 0.5% sodium deoxycholate, 0.1 % SDS, 25 mM β-glycerophosphate, 1 mM EDTA, and 1 mM PMSF) on ice for 15 minutes. The lysate was clarified by centrifugation at 12,000g at 4℃ for 15 minutes. The supernatant was combined with a 1:5 (v/v) ratio of 5×SDS-loading buffer and incubated at 95℃ for 10 minutes. For detecting the spike packaging efficiency, the PSV-containing supernatant was concentrated with a 30% sucrose cushion (30% sucrose, 15 mM Tris-HCl, 100 mM NaCl, 0.5 mM EDTA) at 20,000×g for 1.5 hours at 4℃. The concentrated virus pellet was resuspended in 1×SDS loading buffer and incubated at 95℃ for 30 minutes. For detecting the S2’ cleavage site of PSV, the concentrated viruses were resuspended in DMEM in the presence of indicated concentrations of scFv-mFc or soluble ACE2 for 2 hours at 4 ℃. Then were treated with 10 μg/ml TPCK-trypsin for 30 minutes at 37℃, followed by mixing with a 1:5 (v/v) ratio 5×SDS-loading buffer and incubated at 95℃ for 10 minutes.

After SDS-PAGE and PVDF membrane transfer, blots were blocked with 5% milk in PBS containing 0.1% Tween-20 (PBST) at room temperature for 1 h. Primary antibodies targeting FLAG tag (Sigma-Aldrich, F1804), HA (BioLegend, 901515), VSV-M [23H12] (Kerafast, EB0011), β-tubulin (Immunoway, YM3030) and glyceraldehyde-3-phosphate dehydrogenase (GAPDH) (AntGene, ANT325) were applied at concentrations ranging from 1:2000-1:10,000 in PBST with 1% milk overnight at 4 ℃. The stem-helix targeting monoclonal antibody S2P6 for coronavirus spike detection was used at 1 μg/ml. After three washes with PBST, the blots were incubated with horseradish peroxidase (HRP)-conjugated secondary antibodies (1:10,000). After extensive wash, blots were visualized using the LI-COR Odyssey CLx or the Omni-ECL Femto Light Chemiluminescence Kit (EpiZyme, SQ201) and a ChemiDoc MP Imaging System (Bio-Rad).

### Live-cell binding assays

For detecting coronavirus antigens binding to cell surface expressed CVRs, NTD/CTD/S1-Fc fusion proteins were diluted in DMEM and incubated with cells at the indicated concentrations for 1 h at 37℃. Cells were washed twice with Hanks’ Balanced Salt solution (HBSS) and incubated with 2 μg/mL Alexa Fluor 594 or 488-conjugated goat anti-mouse IgG (Thermo Fisher Scientific; A32742/A32723) for visualization. For detecting the Twin-Strep Tag II labeled S-trimer or soluble ACE2 binding, the incubated cells were treated with 1 μg/mL anti-Twin-Strep Tag II monoclonal antibody (Abbkine; ABT2230) for 30 minutes at 4℃, washed twice with HBSS, and then subjected to fluorescence-labeled secondary antibody incubation. Finally, cells were incubated with Hoechst 33342 (1:5,000 dilution in HBSS) for nuclear staining before imaging using a fluorescence microscope (MI52-N).

### Immunofluorescence assays

Immunofluorescence assays were performed to assess the expression of the CVRs or other receptors carrying the C-terminal 3×FLAG tags. In general, cells expressing the proteins were fixed with 100% methanol at room temperature for 10 minutes, washed once with PBS, and incubated with a mouse monoclonal antibody [M2] specific to the FLAG-tag (Sigma-Aldrich, F1804) in 1% BSA/PBS at 37℃ for 1 hour. After another wash with PBS, cells were incubated with 2 μg/mL Alexa Fluor 594-conjugated goat anti-mouse IgG (Thermo Fisher Scientific, A32742) diluted in 1% BSA/PBS for 1 hour at 37℃. Nuclei were stained with Hoechst 33342 (1:5,000 dilution in PBS). Images were captured and merged using a fluorescence microscope (Mshot, MI52-N).

### Biolayer interferometry assays

Protein binding kinetics were evaluated through Bio-Layer Interferometry (BLI) assays conducted on the Octet RED96 instrument (Molecular Devices). Briefly, 20 μg/mL of S1/NTD/CTD-hFc recombinant proteins were immobilized on protein A (ProA) biosensors (ForteBio, 18-5010). Subsequently, the biosensors were washed and incubated with 2-fold serial-diluted nanobodies (Twin-Strep Tag II) in the kinetic buffer (PBST) to record the association kinetics, followed by recording the dissociation kinetics in the same Kinetic buffer. The background was established using a kinetic buffer without the binding proteins. The kinetic parameters and binding affinities were determined using the Octet Data Analysis software (v.12.2.0.20) through the curve-fitting kinetic analysis or steady-state analysis with global fitting.

### Pseudovirus entry and propagation assays

Single-round VSV-based pseudoviruses carrying the coronavirus spikes were produced following a modified version of a well-established protocol^63^. The VSV-ΔG carrying GFP and firefly luciferase (VSV-ΔG-GFP-fLuc) was rescued using a reverse genetics system in our laboratory, along with helper plasmids from Karafast. For packaging coronavirus PSV, HEK293T cells were transfected with plasmids overexpressing the spike proteins. At 24-36 hours post-transfection, cells were inoculated with 1×10^6^ TCID_50_/mL VSV-dG-GFP-fLuc for 4 hours at 37℃ with 8 μg/mL polybrene. Following two DMEM washes, the culture medium was replenished with DMEM containing 1 μg/mL anti-VSV-G neutralizing antibody (from the I1-mouse hybridoma) to minimize background signals from parental viruses. The TCID_50_ of the PSV was calculated using the Reed-Muench method.

For the pseudovirus propagation assays, the replicable PSVs carrying the GFP reporter and the genomically encoded HKU3 or HKU5 spikes (pVSV-ΔG-GFP-HKU3-S and pVSV-ΔG-GFP-HKU5-S) were generated by the VSV based reverse genetics system. The vector for the VSV genomes was modified based on pVSV-ΔG-GFP-fLuc, with fLuc replaced by the S genes. In brief, the BHK-21 cells were infected with a recombinant vaccinia virus expressing T7 RNA polymerase (vvT7) for 45 minutes at 37℃ (MOI=5). After removing vvT7, the cells were transfected with plasmids containing the pVSV-ΔG-GFP-HKU5/HKU3-S vector and helper plasmids from Karafast. The virus-containing supernatant (P0) was collected 48 hours post-transfection and amplified in Vero E6 cells with in-trans provided VSV-G to yield P1 viruses. The P1 viruses were further amplified in Caco2-CVRs cells in a VSV-G independent manner and in the presence of anti-VSVG (I1-Hybridoma supernatant), generating P2 viruses that were dependent on the genomically encoded HKU3 and HKU5 spike proteins for amplification.

For pseudovirus entry or entry inhibition assays, susceptible cells were cultured in 96-well plates at a density of 5×10^4^ cells per well and then incubated with around 1×10^6^ TCID_50_/mL of pseudovirus (PSV), with 100 μL per well. The incubation allowed for attachment and viral entry with or without the indicated concentrations of antibodies or other inhibitors. In some cases, TPCK-treated trypsin of indicated concentrations (sigma, T8802) was added to the medium to enhance entry efficiency. At 16-20 hour post infection (hpi), 40 μL of One-Glo-EX substrate (Promega) was added to the cells and incubated for at least 5 minutes on a plate shaker in the dark. Relative light units (RLU) were determined using the GloMax 20/20 Luminometer (Promega). GFP intensity was analyzed using a fluorescence microscope (Mshot, MI52-N).

### Cell-cell fusion assays

A cell-cell fusion assay based on dual split proteins (DSPs) was performed on HEK293T or BHK21-T7 cells stably expressing the CVRs or the native receptors^16^. Group A cells were transfected with plasmids expressing spike protein and RLucN(1-155)-sfGFP(1-157), while the group B cells were transfection with plasmids expressing spike proteins (same as in group A) and sfGFP(158-231)-RLuc(156-311). Cells from both groups were trypsinized and co-cultured in a 96-well plate at a density of approximately 1×10^5^ cells per well at 12 hours post-transfection. After 16-24 hours, cell nuclei were stained with Hoechst 33342 (1:5,000 dilution in HBSS) for 30 minutes at 37 ℃, and the fluorescent images were captured using a fluorescence microscope (MI52-N; Mshot). For the assessment of live-cell luciferase activity after reconstitution of split RlucN, 20 μM of EnduRen live-cell substrate (Promega, E6481) was added to the cells in DMEM and incubated for at least 1 hour before detection using the Varioskan LUX Multi-well Luminometer (Thermo Fisher Scientific).

### Flow cytometry analysis

For flow cytometry analysis, viral antigen-mFc and VBDs-mFc recombinant proteins were diluted in DMEM at the indicated concentrations and then incubated with HEK293T cells expressing the indicated receptors or coronaviruses spike proteins for 1 hour at 37℃. In live cell binding assays, for detecting the cell surface hFc or intracellular ZsGreen, cells were washed with DMEM and subsequently incubated with either Alexa Fluor 594-conjugated goat anti-mouse IgG (Thermo Fisher Scientific; A32742) or a combination of Alexa Fluor 488-conjugated goat anti-human IgG (Thermo Fisher Scientific; A11013). In live cell binding assays, for detecting the cell surface 76E1 epitope exposure, the SARS2-CoV-2-S IRES-ZsGreen expressing cells were incubated with indicated concentrations of scFv-mFc or soluble receptors for 1 hour at 37℃ before 76E1 antibody incubation (1 μg/mL). When detection of the intracellular FLAG tag is necessary, cells were washed once with HBSS and fixed with 4% PFA, permeabilized with 0.1% Triton X-100, blocked with 1% BSA/PBS at 4℃ for 30 minutes, and subsequently stained with Rabbit anti-Flag tag mAb (CST,14793S) diluted in 1% BSA/PBS for 1 hour at 4℃ to visualize the expression of CVRs and other receptors. Following extensive washing, the cells were incubated with Alexa Fluor 647-conjugated goat anti-rabbit IgG (Thermo Fisher Scientific; A32733) and Alexa Fluor 488-conjugated goat anti-mouse IgG (Thermo Fisher Scientific; A32723), both diluted in 1% BSA/PBS, for 1 hour at 4℃. Following the completion of all staining procedures, cells washed twice with PBS were subsequently analyzed using the CytoFLEX Flow Cytometer (Beckman). In each case, 5,000 cells expressing either receptors or spikes, gated based on FLAG/hFc/ZsGreen-fluorescence intensity and SSC/FSC, were analyzed with the CytoFLEX Flow Cytometer (Beckman).

### Reverse genetics to rescue HKU5-WT and HKU5-ZGH

The full-length cDNA clone of HKU5 (GenBank: NC_009020) was designed and synthesized as seven (from A to G) contiguous cDNAs flanked by unique class IIS restriction endonuclease site (BsaI or BsmBI) and cloned in pUC57 vector. Class II restriction endonuclease sites AvrII and AscI were introduced to 5’ terminal of HKU5 A and 3’ terminal of HKU5 G fragments, respectively. Several silent mutations were included to disrupt naturally occurring restriction cleavage sites. A poly-A (25 repeats) sequence was introduced to 3’ terminal of HKU5 G fragment. To assemble the full-length cDNA clone, HKU5 A-G fragments were digested by endonucleases, resolved on 1% agarose gels, purified with a gel extraction kit, extracted with chloroform, and precipitated with isopropyl alcohol. Digested HKU5 A-G inserts, and modified pBaloBAC11 vector were mixed, ligated overnight at 4℃, and transformed into DH10B competent cells. The correct full-length HKU5 cDNA clone was identified and verified by sequencing. The construction of HKU5-ZGH utilized the transformation-associated recombination (TAR) cloning technique. Specifically, a ZsGreen-HiBit (ZGH) DNA fragment was commercially synthesized (Tsingke) to replace the HKU5-ORF5. The PCC1 vector was used to clone the HKU5 genomic DNA carrying the ZGH substitution based on three segments amplified using the HKU5-WT infectious clone as a temperate. Subsequently, all the products were transformed into yeast using the high-efficiency lithium acetate/SS carrier DNA/PEG method. The yeast plasmid was extracted and transformed into EPI300 electrocompetent cells. The plasmid used for cell transfection was obtained from a 300 mL E. coli bacterial culture suspension. For transfection, 4 μg of both HKU5 WT and HKU5-ZGH plasmids were separately transfected into Caco2-1B4 cells (1×10^6^ cells) using Lipofectamine 2000. Progeny viruses collected from the supernatant at 72 hours post-transfection (P0) were utilized to generate stocks for subsequent analyses.

### Transmission electron microscopy

Viral culture supernatant was fixed with formaldehyde (working concentration 0.1%) at 4℃ overnight. Subsequently, it was concentrated by ultracentrifugation through OptiPrepTM Density Gradient Medium (D1556) at 154,000 g at 4℃ for 2.5 hours using a SW41Ti rotor (Beckman). The pelleted viral particles were suspended in 100 μL of PBS, stained with 2% phosphotungstic acid (pH 7.0), and examined using a Tecnai transmission electron microscope (FEI) at 200 kV.

### Authentic coronavirus infection assays

Human coronavirus 229E (VR-740) is obtained from ATCC and amplified in Huh-7 cells. MHV-A59 is a gift from Professor Yu Chen’s lab (Wuhan University) and is amplified in Neuro2a cells. The SARS-CoV-2-ΔN with N protein substituted with EGFP is rescued using an established protocol, and cultured in Caco2 cells overexpressing the SARS-CoV-2 N protein^64^. All experiments involving RsHuB2019A, HKU5-WT, and HKU5-ZGH authentic viruses infection were conducted in the certified negative-pressure Biosafety Level 2 laboratory at Wuhan Institute of Virology. RsHuB2019A is amplified in either Huh-7 or in Caco2-Nb27 cells. HKU5-WT and HKU5-ZGH are amplified in Caco2-1B4 cells.

For replication experiments, target cells were initially seeded in 24-well plates and washed with DMEM before inoculation, either in the presence or absence of trypsin (100 μg/mL). Following a one-hour incubation at 37℃, the cells were washed with DMEM and further incubated for the indicated hours at 37℃. For qRT-PCR analysis, cell-free supernatants (50 μL per well each time) were collected at indicated time points post-infection and stored at −80℃. Viral RNA was extracted using Virus DNA/RNA Extraction Kit (Vazyme: RM501) and subjected to qRT-PCR as previously described^65^. Primers for RsHuB2019A RdRp: 5’-TTGTTCTTGCTCGCAAACATA-3’ (forward) and 5’-CACACATGACCATCTCACTTAA-3’ (reverse). Primer for HKU5 nsp2: 5’-CTGCGCTTAATGCCCCATTC-3’ (forward) and 5’-GACGTGTAGACGTAGAGCCG-3’ (reverse). Primers for VSV L protein, forward primer: 5’-TCTTGAGTTGTGGAGACGGC-3’ (forward) and 5’-ACCGTCTTGAACATGGGACC-3’ (reverse). Primers for MHV-A59 N protein:5’-TATAAGAGTGATTGGCGTCC-3’(forward) and 5’-GAGTAATGGGGAACCACACT-3’ (reverse). All samples were analyzed in duplicate on two independent runs.

For immunofluorescence assays, cells were fixed with methanol for 40 minutes at room temperature at indicated time points. The expression of RsHuB2019A and HKU5 N proteins was detected by rabbit anti-SARS-related CoV Rp3 N protein serum (diluted at 1:2000) and rabbit anti-HKU5 N protein serum (diluted at 1:4000), respectively, followed by DL594-conjugated goat anti-rabbit IgG (Thermo, 1:1000) staining. MHV-A59 and 229E-VR740 spike proteins were detected using their Spike-targeting nanobodies (1A1-mFc for 229E-VR740; 1F7-mFc for MHV-A59), followed by DL594-conjugated goat anti-mouse IgG antibodies (Thermo, 1:1000).

### Nanobody bio-panning

Specific viral antigens (30-100 μg) were immobilized on streptavidin-conjugated magnetic beads for one-hour incubation at 37℃ and extensively washed to remove unbound antigens. Subsequently, the beads were incubated with the nanobody library (1×10^10^ PFU) (Naïve VHH libraries from Camelus bactrianus, Alpaca, and Llama from NBbiolab, China) for 1 hour. The bound phages were eluted using an Elution Buffer (50 mM Tris-pH 7.4, 150 mM NaCl, 50 mM biotin) after extensive washing with PBST to eliminate nonspecific binders. The eluted phage encoding the specific nanobodies was proliferated in E. coli (TG1). After one round of magnetic beads-based selection. 1-3 additional rounds of phage biopanning were conducted using magnetic beads or immunotubes. The positive clones were identified through the enzyme-linked immunosorbent assays (ELISA), sequenced, and verified by cell-based binding assays.

### Biosafety and biosecurity

Experiments related to authentic human coronavirus 229E, SARS-CoV-2-ΔN, and murine MHV-A59 were authorized by the Biosafety Committee of the State Key Laboratory of Virology, Wuhan University and conducted in accordance with standard operating procedures (SOPs) in a BSL-2 laboratory. SARS-CoV-2 authentic viruses-related experiments were conducted in the ABSL-3 facility at Wuhan University with the approval of the Biosafety Committee of the ABSL-3 laboratory. The SARS-CoV-2 WT strain (IVCAS 6.7512) was provided by the National Virus Resource, Wuhan Institute of Virology, Chinese Academy of Sciences and amplified in Vero E6 cells in the ABSL-3 facility at Wuhan University. Experiments related to authentic viruses RsHuB2019A, HKU5-WT, and HKU5-ZGH were approved by the Wuhan Institute of Virology (WIV) IBCs and performed in the BSL-2 laboratory according to SOPs at WIV facilities. All the facilities at both Wuhan University and WIV for this work adhere strictly to the safety requirements recommended by the China National Accreditation Service for Conformity Assessment.

### Statistical analysis

Most experiments were repeated 2-5 times, each with approximately 3-4 biological replicates. Results are presented as mean ± standard deviation (s.d.) or mean ± standard error of the mean (s.e.m.), as specified in the figure legends. Statistical analyses were primarily performed using GraphPad Prism (V.8) through unpaired two-tailed Student’s t-tests two independent groups or One-way ANOVA analysis followed by Dunnett’ s test for multiple comparisons. P < 0.05 was considered statistically significant; *P < 0.05; **P < 0.01; ***P < 0.001; ****P < 0.0001; ns, non-significant.

---

**Extended Data Fig. 1.**
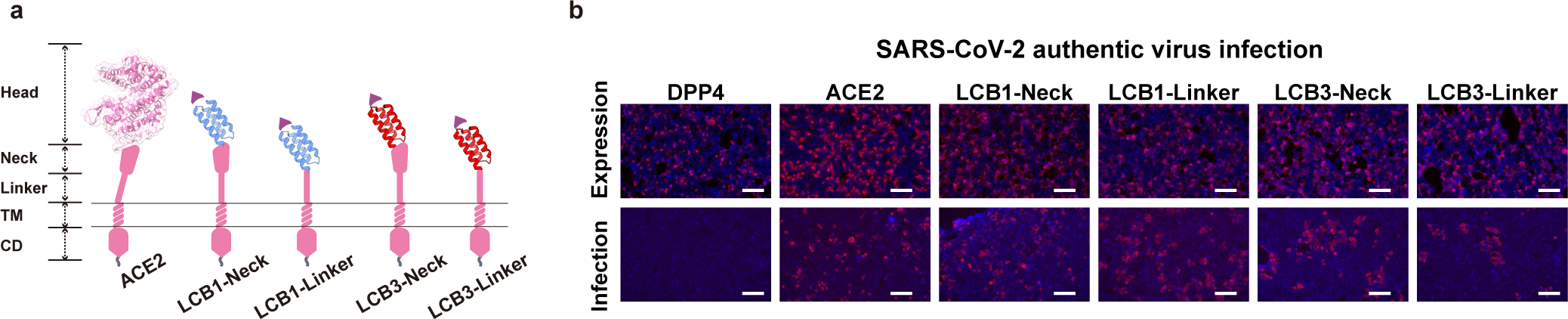
Efficient authentic SARS-CoV-2 infection supported by chimeric ACE2 with viral binding domain substituted by ACE2-mimicking helical frameworks (Hf). **a,** Schematic illustration illustrating the four Hf-based CVRs. **b**, Immunofluorescence analysis of ACE2 or CVRs expression levels stably expressed in HEK293T cells, detected by C-terminal fused 3×FLAG tags. **c**, Immunofluorescence analysis of authentic SARS-CoV-2 infection efficiencies in indicated cells by detecting intracellular SARS-CoV-2 N proteins at 24 hour post infection (hpi). Scar bars:100 μm.

**Extended Data Fig. 2.**
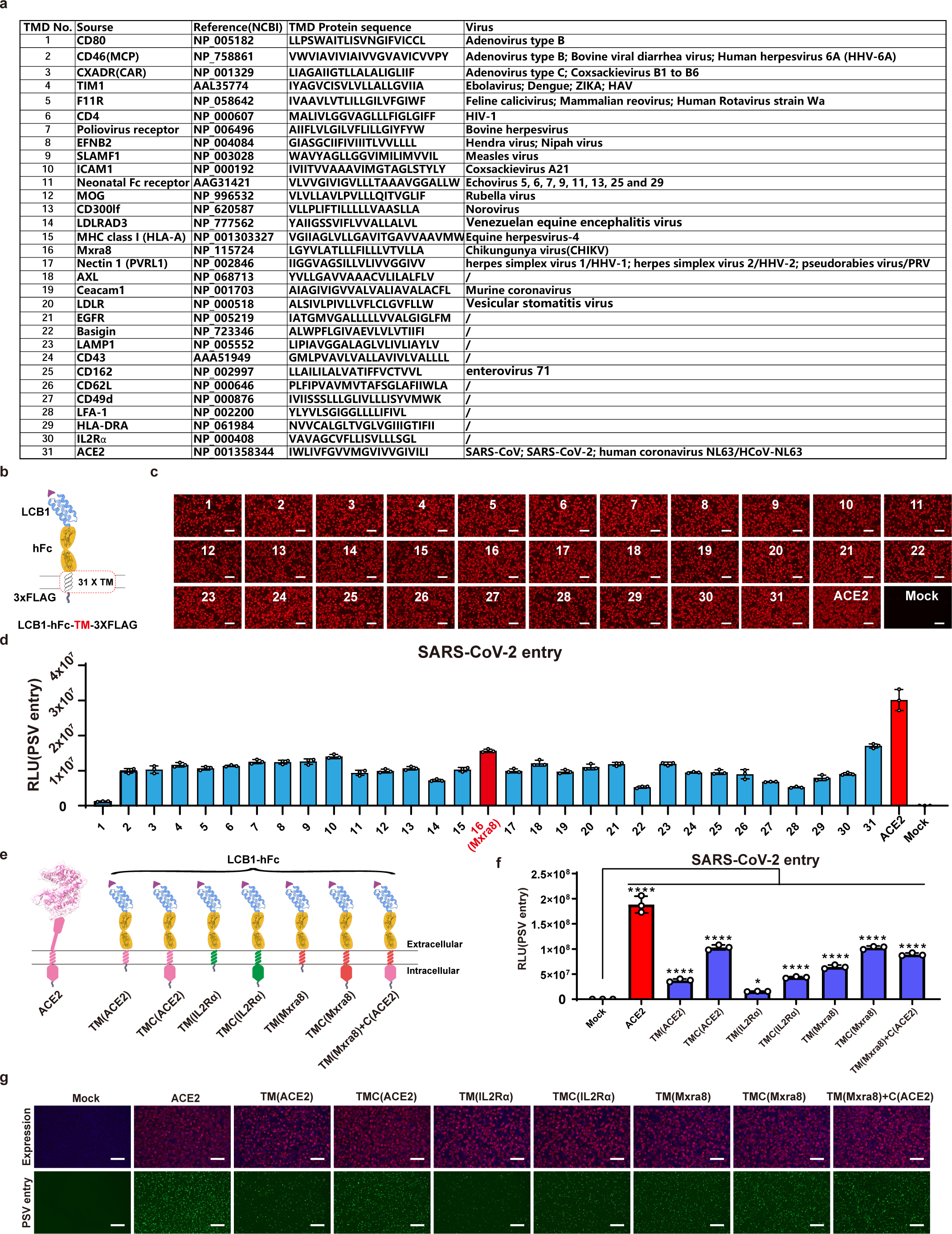
Comparison of CVRs carrying transmembrane and cytosolic domains from different receptors. **a,** Details of the 31 different TM sequences examined in this study. **b,** Cartoon illustrating the framework of the CVRs for TM evaluation. **c**, Immunofluorescence analysis of the expression of the 31 CVRs in HEK293T cells by detecting the C-terminal fused 3× FLAG tags. **d**, Evaluation of SARS-CoV-2 PSV entry efficiency supported by the indicated CVRs carrying different TMs. Mxra8 TM displaying the best performance was marked in red. **e**, Cartoon illustrating the LCB1-based CVRs with selected TM or TMC substitutions for further verification. **f, g,** PSV entry-supporting efficiencies of the CVRs assessed by RLU(**f**) or GFP reporters (**g**) in transiently transfected 293T cells. Scare bars: 100 μm. One-way ANOVA analysis followed by Dunnett’s test for **f.**

**Extended Data Fig. 3.**
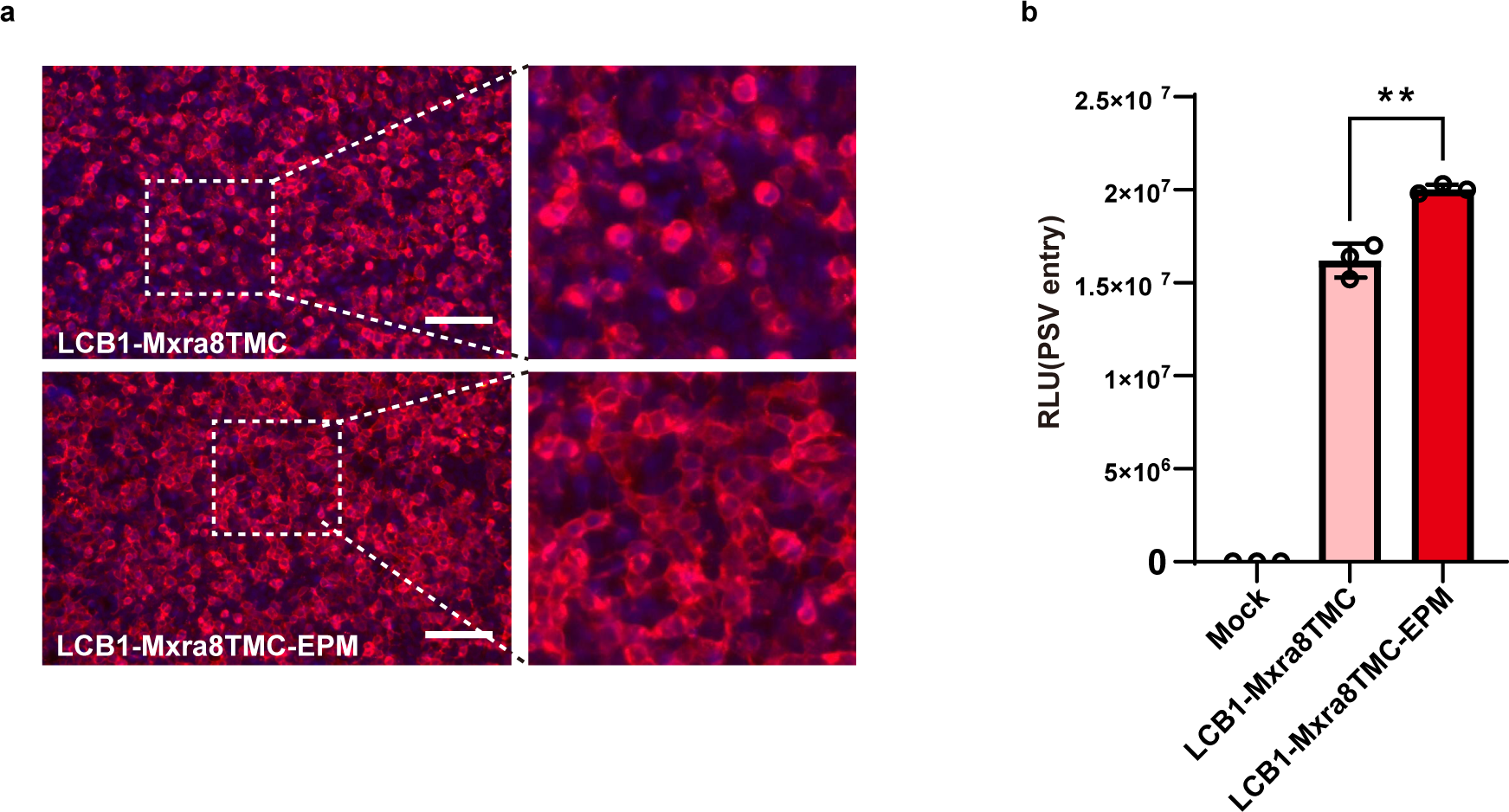
EPM promotes cell surface distribution and entry-supporting efficiency of CVRs. **a,** Immunofluorescence displaying the subcellular distribution of LCB1-Mxra8TMC-based CVRs transiently expressed in HEK293T cells with or without EPM. The white dashed boxes highlight the cell surface distribution at a higher magnification. **b,** Evaluation of the SARS-CoV-2 PSV entry efficiency of the CVRs with or without the EPM. Scare bars: 100 μm. Unpaired two-tailed Student’s t-tests for **b**.

**Extended Data Fig. 4.**
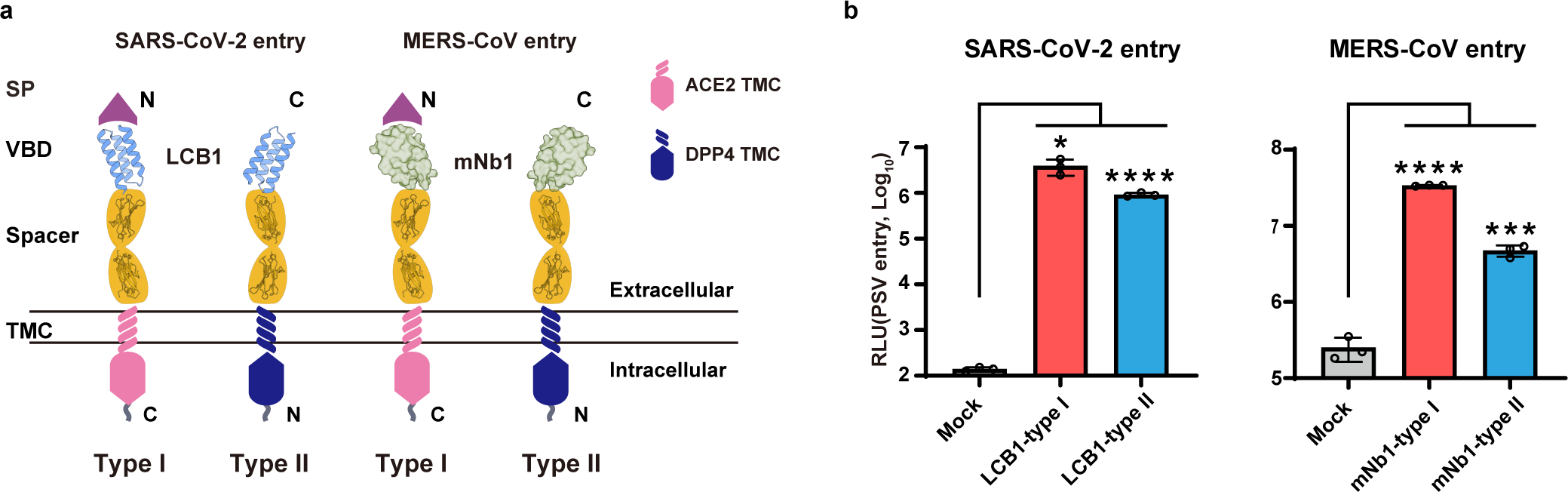
Comparison of CVRs displayed with different transmembrane topologies. **a,** Cartoon illustrating CVRs carrying LCB1 or mNb1 displayed in either type I or type II transmembrane topology. **b,** Evaluation of SARS-CoV-2 or MERS-CoV PSV entry efficiency supported by the indicated CVRs with different transmembrane topologies in HEK293T cells. Unpaired two-tailed Student’s t-tests for **b**.

**Extended Data Fig. 5.**
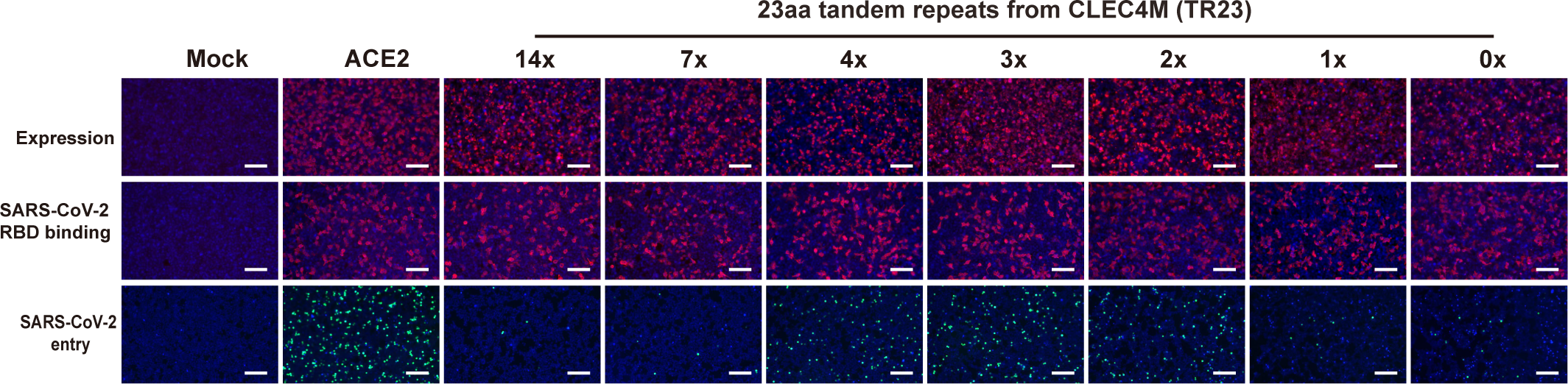
Comparison of CVRs carrying varying copies of TR23 repeats as spacers. Assessment of CVR expression, SARS-CoV-2 RBD-mFc binding, and PSV entry efficiency supported by the indicated CVRs transiently expressed in HEK293T cells. Scare bars: 100μm.

**Extended Data Fig. 6.**
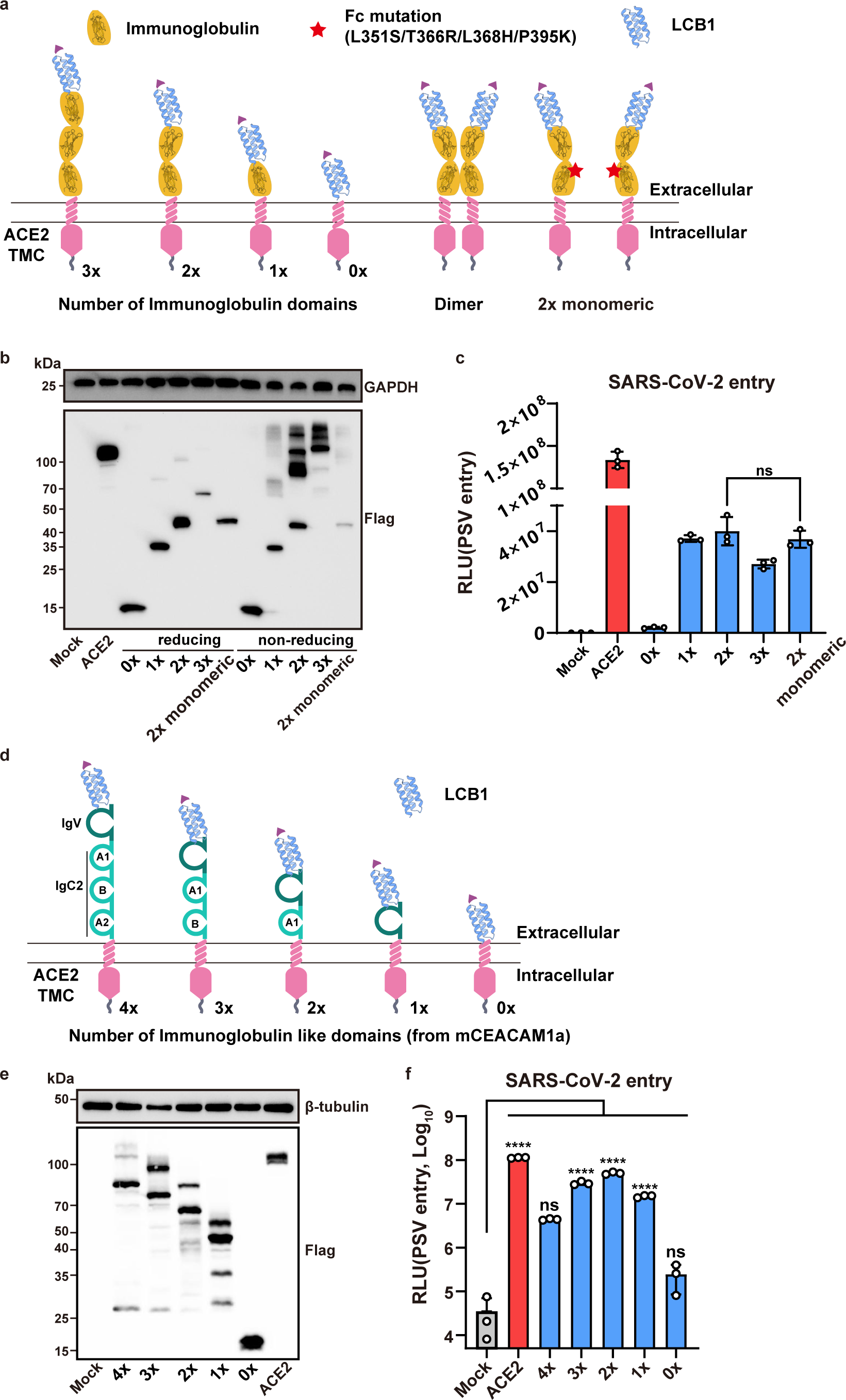
Comparison of CVRs carrying different numbers of immunoglobulin domains or dimerization-abolished hFc as spacers. **a,** Schematic representation of the CVRs carrying different numbers of immunoglobulin (Ig) domains (left) or an Fc mutant with abolished dimerization ability. **b,** Western blot analysis of CVRs expression in HEK293T cells under either reducing or non-reducing conditions, respectively. **c,** Assessment of SARS-CoV-2 PSV entry efficiency in HEK293T cells transiently expressing the indicated CVRs. **d,** Schematic representation of the CVRs carrying different numbers of Ig-like domains (left) from mCEACAM1a. **e,** Western blot analysis of CVRs expression in HEK293T cells. **f,** SARS-CoV-2 PSV entry efficiency in HEK293T cells transiently expressing the indicated CVRs. Unpaired two-tailed Student’s t-tests for **c**. One-way ANOVA analysis followed by Dunnett’s test for **f**.

**Extended Data Fig. 7.**
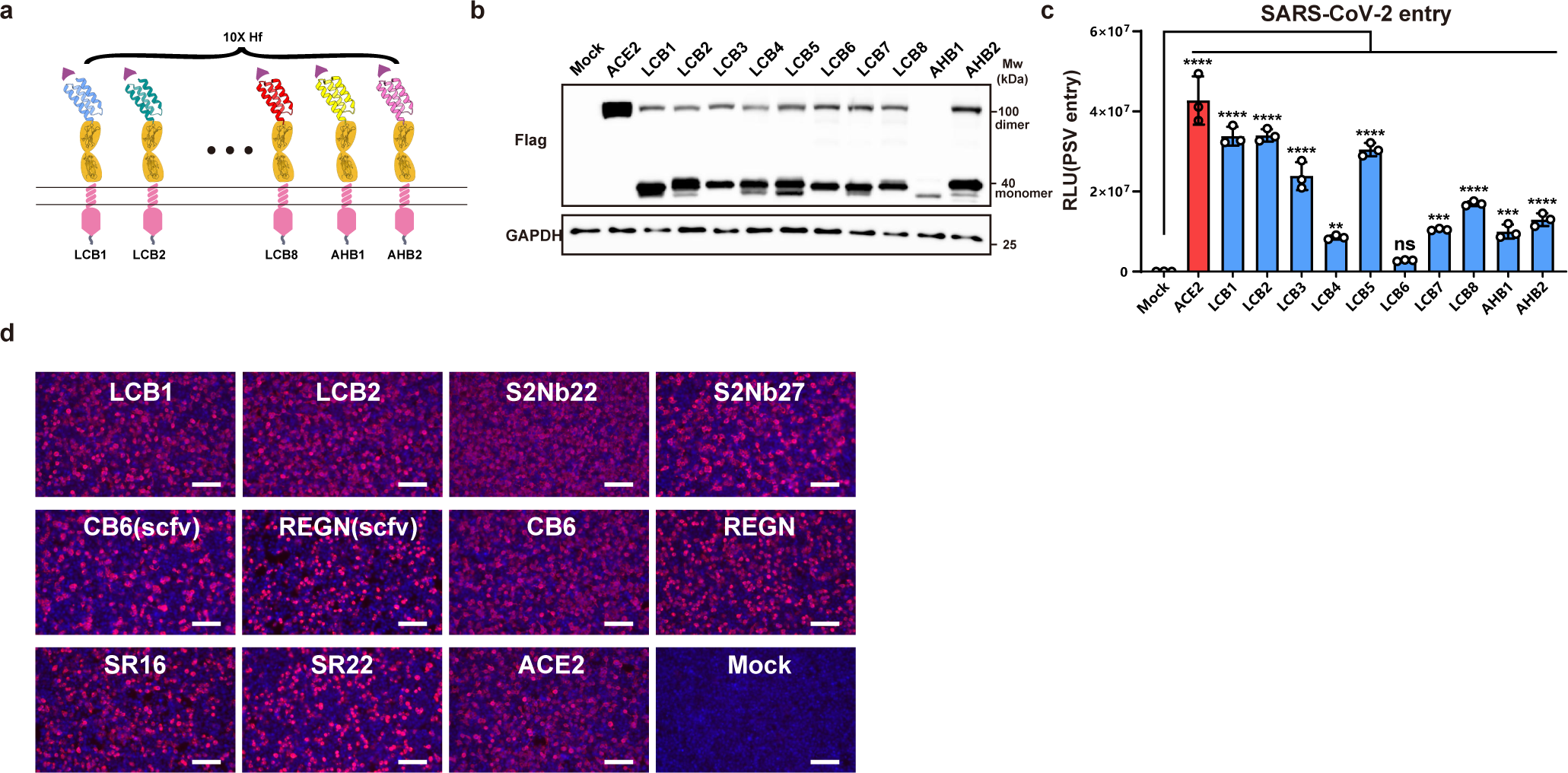
Expression levels and entry-supporting efficiency of CVRs carrying different viral binding domains. **a**, Schematic representation of the CVRs carrying different ACE2-mimicking Hf. **b, c**, Expression (b) and SARS-CoV-2 entry-supporting (c) ability of different CVRs in 293T cells. **d,** Immunofluorescence analyzing the expression of the indicated CVRs transiently expressed in HEK293T cells by detecting the C-terminal fused 3×FLAG tags. Scare bars: 100μm. One-way ANOVA analysis followed by Dunnett’s test for **c**.

**Extended Data Fig. 8.**
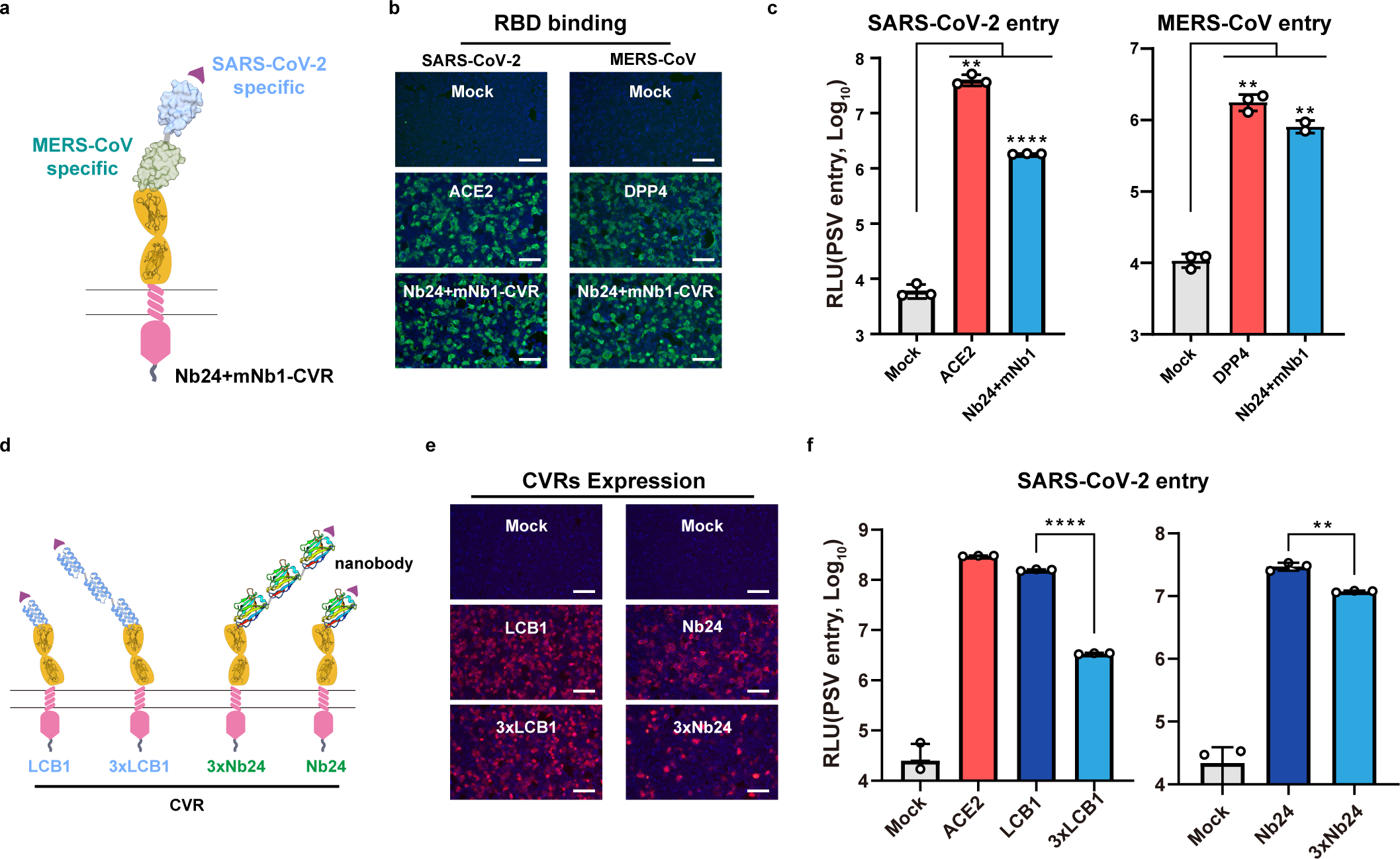
Functionality of CVRs carrying bi-specific VBDs or polymeric VBDs in supporting coronavirus entry. **a-c,** Illustration (a), viral RBD binding efficiency (b), and PSV entry-supporting efficiency (c) of a SARS-CoV-2/MERS-CoV bi-specific CVR transiently expressed in HEK293T cells. **d-f,** Illustration (d), expression (b), and PSV entry-supporting efficiencies (c) of CVRs carrying a single VBD or tandemly connected VBD trimmer. Scare bars: 100μm. Unpaired two-tailed Student’s t-tests for **c,** and **f**.

**Extended Data Fig. 9.**
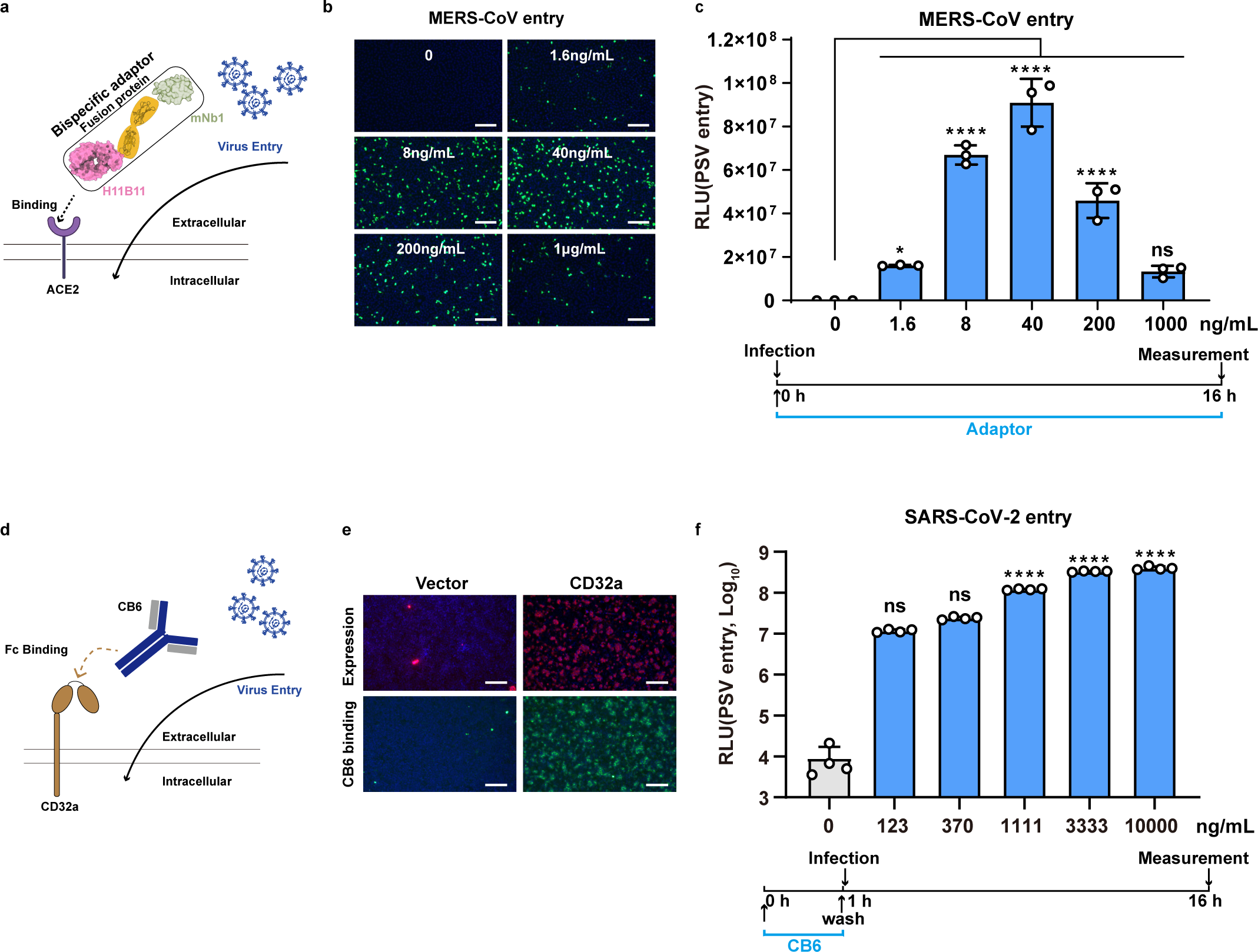
Coronavirus tropism retargeting mediated by bi-specific soluble adaptor proteins. **a-c,** Schematic illustration of bispecific adaptor protein (**a**) and MERS-CoV PSV entry efficiency in BHK-21-hACE2 cells in the presence of indicated concentrations of adaptor proteins (H11B11-mNB1) throughout the infection. PSV entry efficiency is examined based on the GFP intensity (**b**) or RLU (**c**) in the infected cells. **d-f**, Schematic illustration of FcγR (CD32a) mediated antibody-dependent coronavirus entry (**d**). CD32a expression, antibody (CB6) binding (**e**), and SARS-CoV-2 PSV entry (**f**) into HEK293T-CD32 cells, which was pretreated with indicated concentration (con.) of the CB6 for 0.5h. Scare bars: 100 μm. One-way ANOVA analysis followed by Dunnett’s test for **c** and **f**.

**Extended Data Fig. 10.**
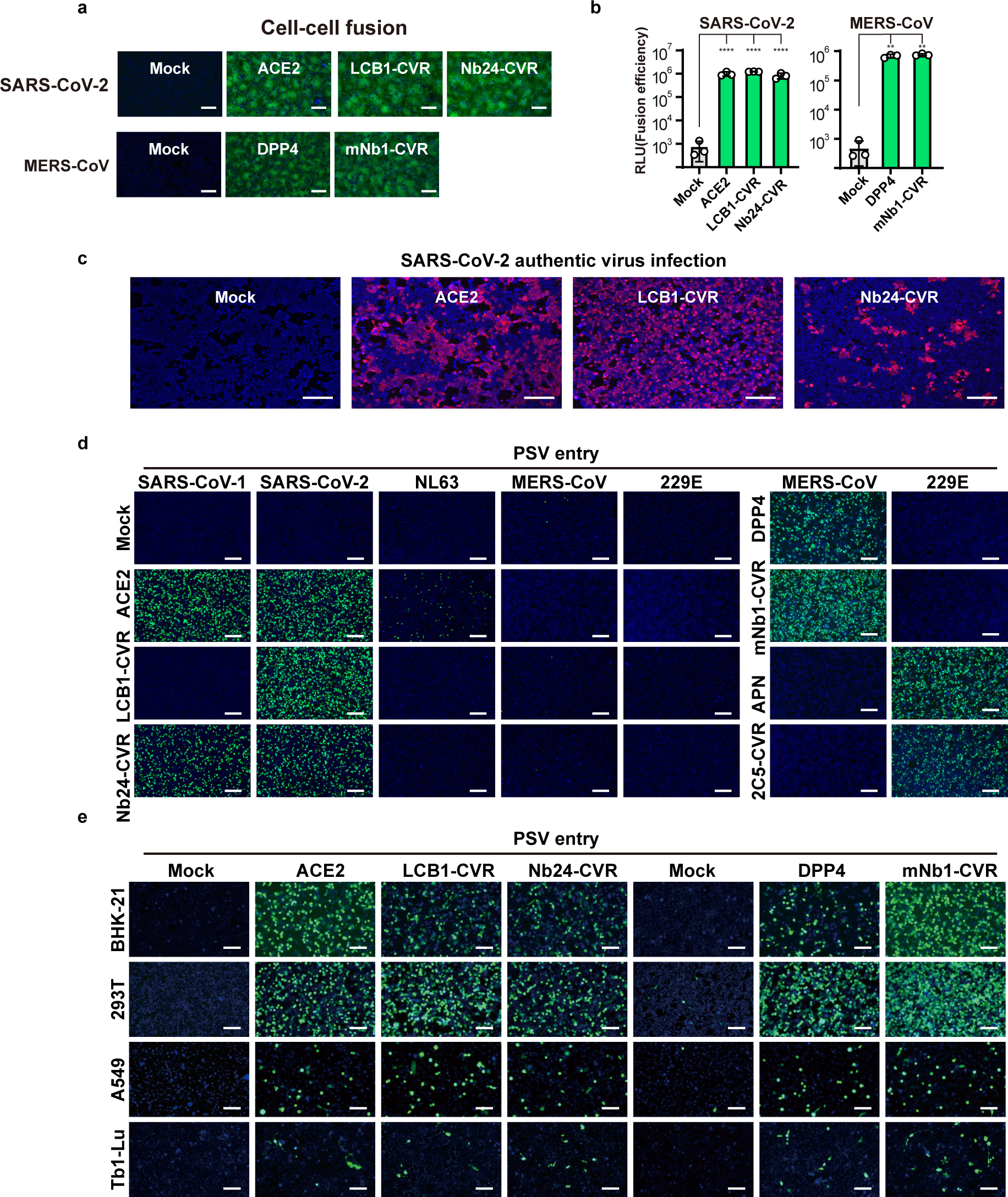
Comparison of specificity and receptor functionality between CVRs and native receptors in different cell types. **a-c,** Evaluation of the ability of CVRs to induce cell-cell fusion (**a,b**) and to support SARS-CoV-2 authentic virus infection (**c)**. Spike-receptor mediated cell-cell fusion was demonstrated by the reconstituted GFP (a) and Renilla luciferase activity (RLU) (**b**). Infection was analyzed by immunofluorescence detecting the intracellular N protein at 24 hpi (**c**). **d,** Entry of different coronavirus PSVs into HEK293T stably expressing the native receptor or the indicated CVRs. **e,** SARS-CoV-2 and MERS-CoV PSV entry into various cell types expressing the indicated receptors. Scare bars: 100 μm. Unpaired two-tailed Student’s t-tests for **b**.

**Extended Data Fig. 11.**
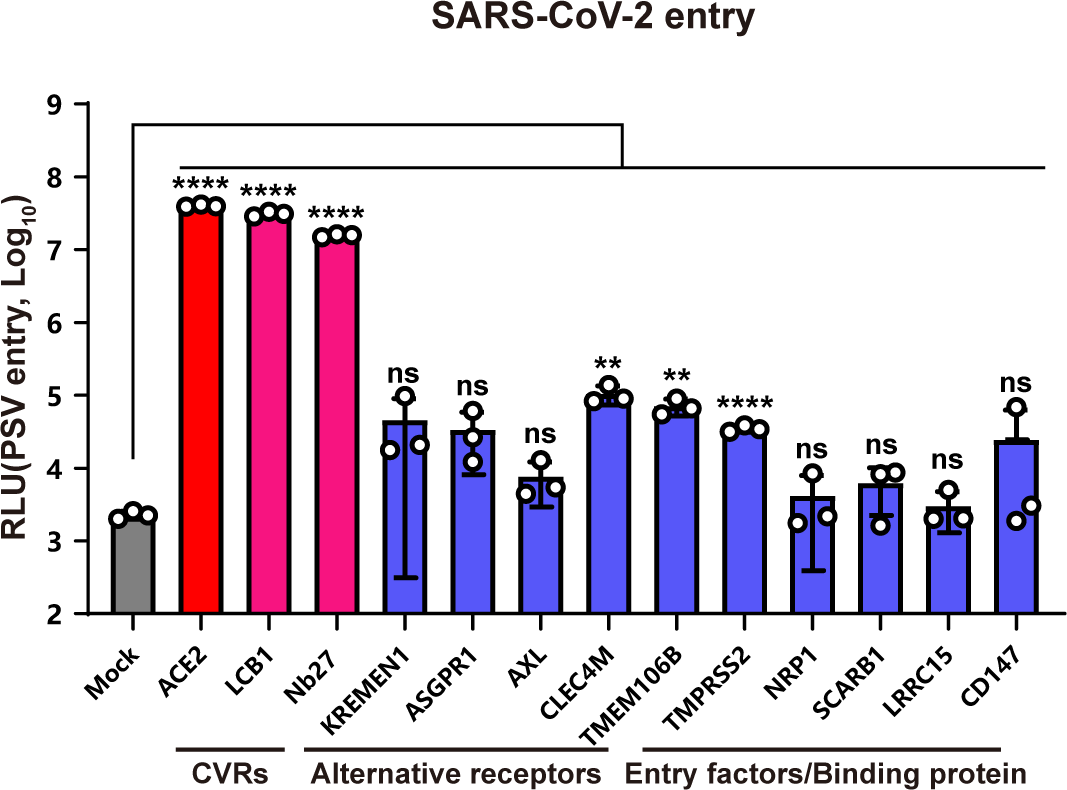
Comparison of the SARS-CoV-2 entry efficiency supported by ACE2, CVRs, alternative receptors, or other entry factors. SARS-CoV-2 PSV entry in HEK293T cells expressing the indicated receptors or entry factors. Unpaired two-tailed Student’s t-tests was employed for analysis.

**Extended Data Fig. 12.**
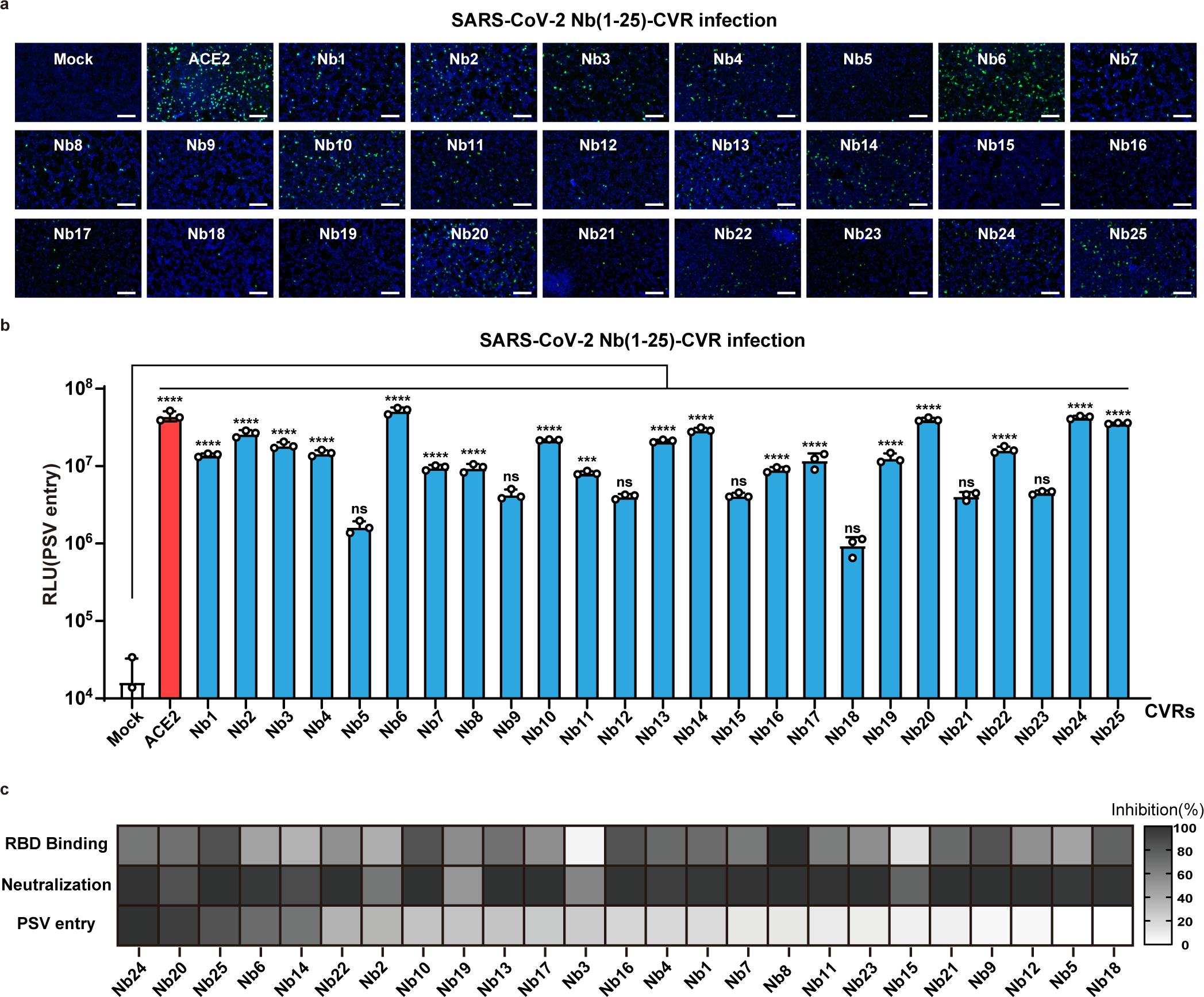
Relationship between the antigen binding, soluble antibody neutralizing activity, and CVR entry-supporting ability of 25 SARS-CoV-2 RBD targeting nanobodies. **a, b,** Assessment of the entry-supporting ability of 25 nanobody-CVRs in HEK293T cells, indicated by GFP(**a**) and the RLU (**b**), respectively. **c,** Comparison of RBD-mFc binding, soluble nanobody-hFc neutralization, and PSV entry efficiencies in HEK293T cells. RBD-mFc binding and PSV entry assays were conducted in HEK293T transiently expressing the 25 CVRs. The SARS-CoV-2 PSV neutralization assay was performed in HEK293T-ACE2 in the presence of indicated nanobody-Fc recombinant proteins. Scare bars: 100μm. One-way ANOVA analysis followed by Dunnett’s test for **b**.

**Extended Data Fig. 13.**
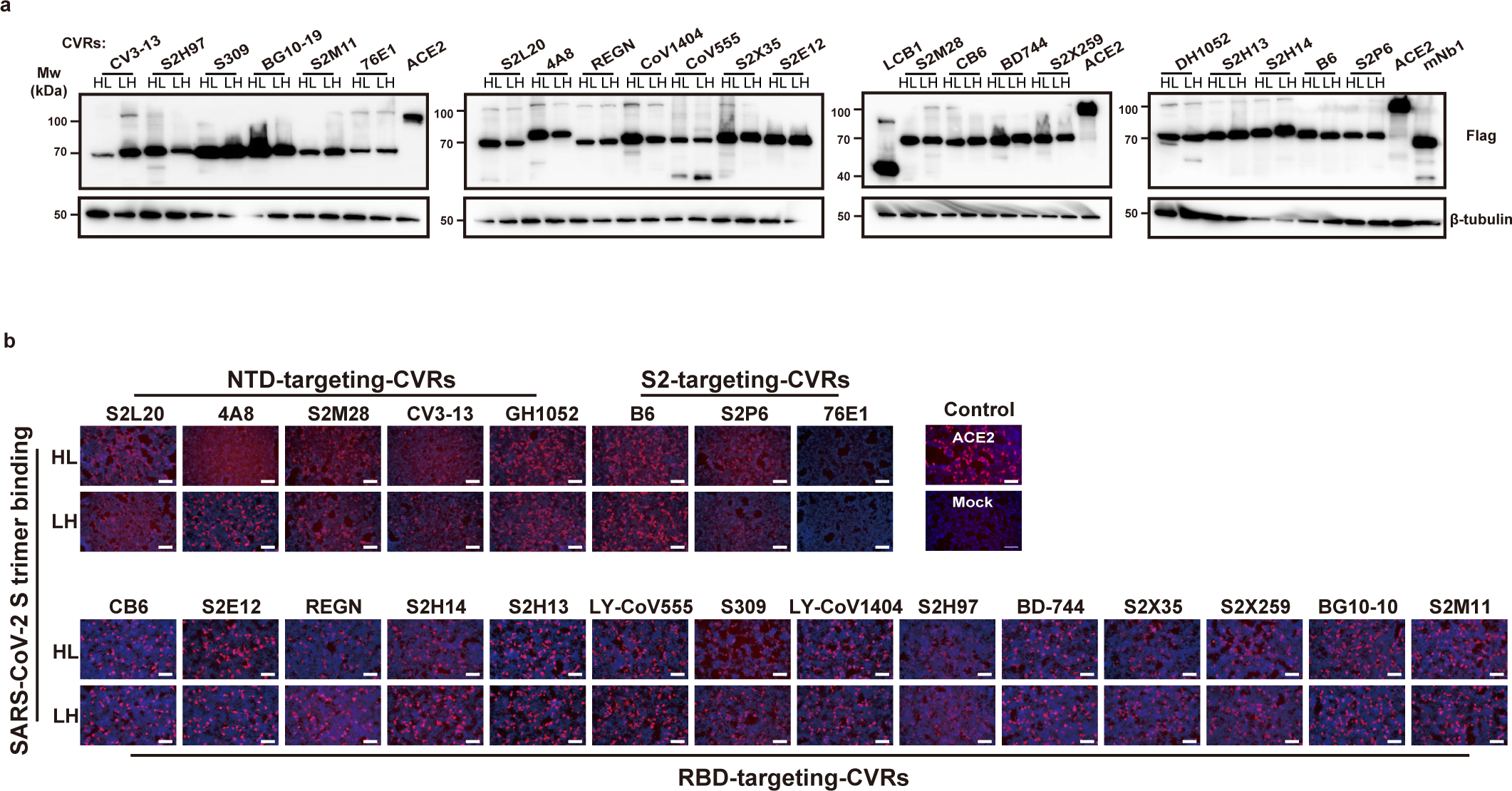
Expression and SARS-CoV-2 spike trimmer binding efficiencies in cells expressing the indicated scFv-CVRs. **a,** Western blot analysis of the expression levels of indicated scFv-CVRs transiently expressed in HEK293T cells. **b**, Binding of SARS-CoV-2 S-trimer to HEK293T cells expressing the indicated CVRs.

**Extended Data Fig. 14.**
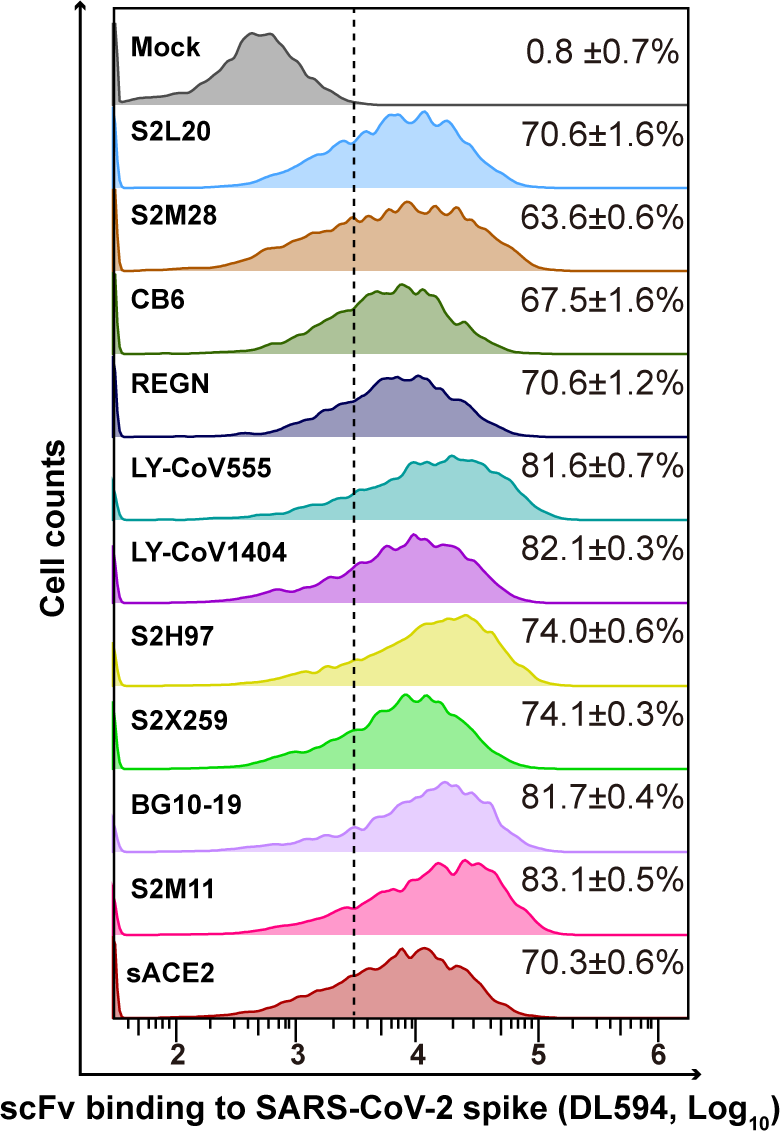
Binding efficiencies of scFv-mFc targeting different SARS2-CoV-2 epitopes in cells expressing the SARS-CoV-2 spike. Flow cytometry analysis was performed to assess the binding efficiency of scFv-mFc with HEK293T cells transiently expressing the SARS-CoV-2 Spike proteins and ZsGreen simultaneously. The ZsGreen positive cells were gated for subsequent analysis of mFc binding efficiency.

**Extended Data Fig. 15.**
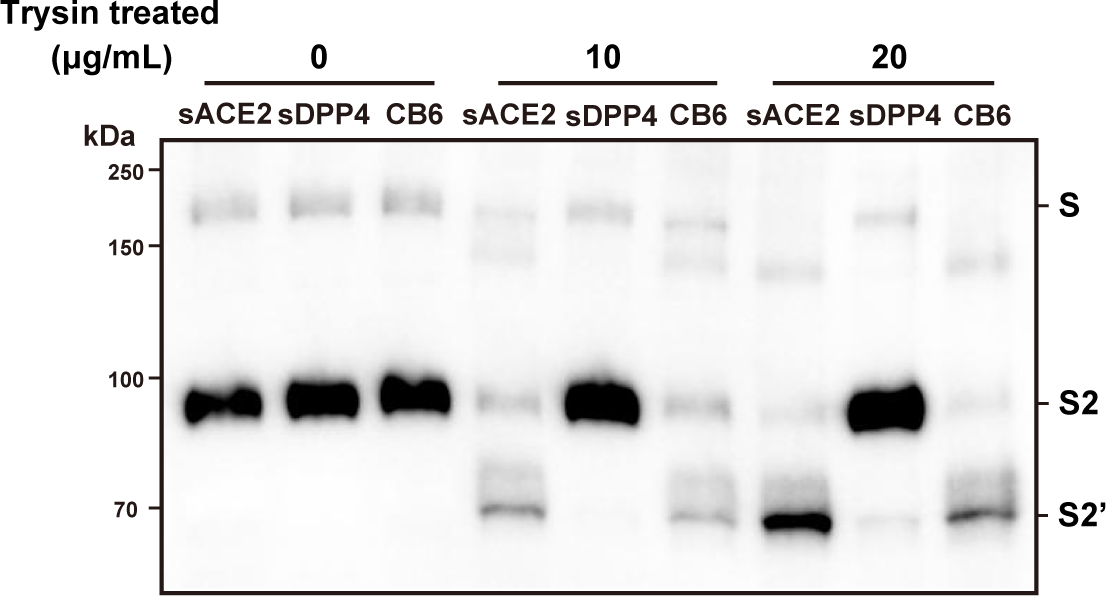
Trypsin-mediated S2’ cleavage of SARS2-CoV-2 PSV in the presence of soluble receptors or CB6-scFv-mFc. The concentrated SARS-CoV-2 PSV particles were incubated with 100 μg/mL of soluble receptors or CB6-scFv-mFc for 1 hour, followed by incubation with the indicated concentration of TPCK-treated trypsin for 30mins. Western blot analysis was conducted by detecting the S2P6 epitope on the S2 subunit. sDPP4: soluble DPP4 ectodomain proteins.

**Extended Data Fig. 16.**
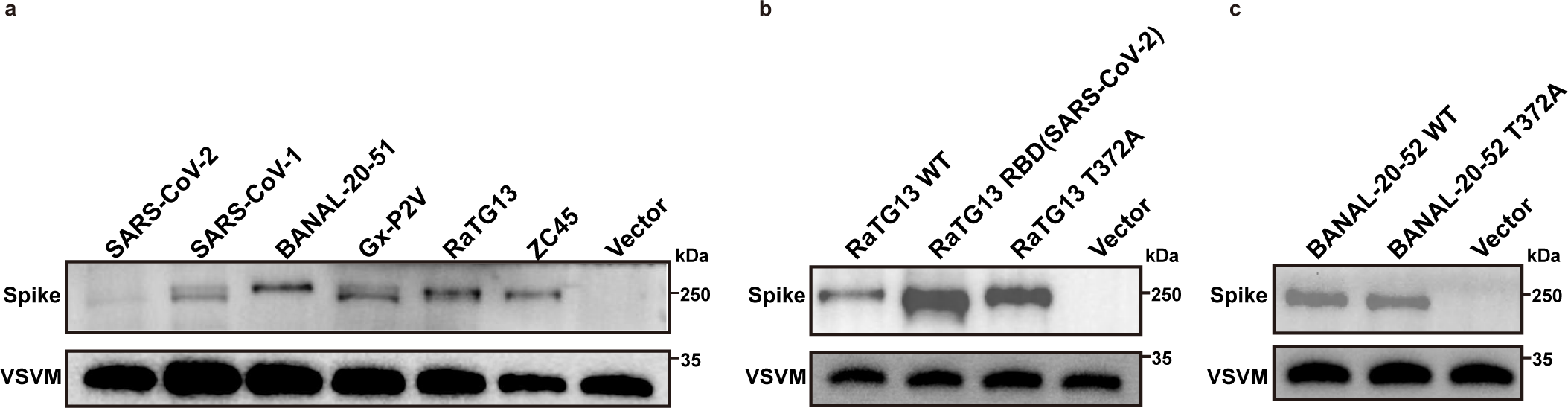
Package efficiency of PSVs carrying indicated coronavirus spike proteins. Western blot detection of concentrated PSV carrying indicated coronaviruses by detecting the S2P6 epitope conserved among the tested coronaviruses. VSV-M serves as a loading control.

**Extended Data Fig. 17.**
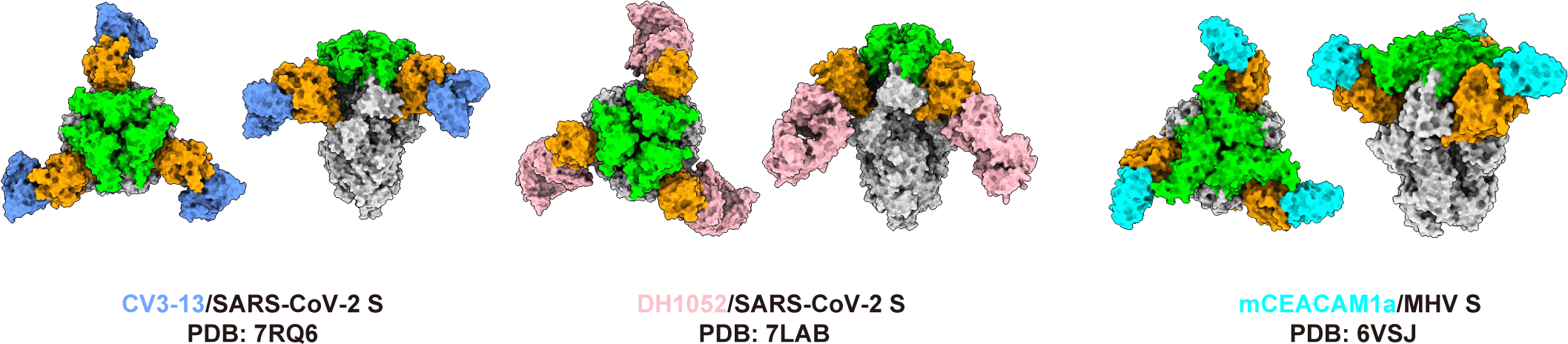
Cryo-EM structures of NTD-targeting antibodies or soluble mCEACAM1a in complex with SARS-CoV-2 or MHV spike trimmer, respectively. Illustration of top-view and side-view cryo-EM structures depicting NTD-targeting antibodies (CV3-13 and DH1052) or soluble mCEACAM1a in complex with SARS-CoV-2 or MHV spike trimmer, respectively. The complex structures are annotated with corresponding PDB accession numbers.

**Extended Data Fig. 18.**
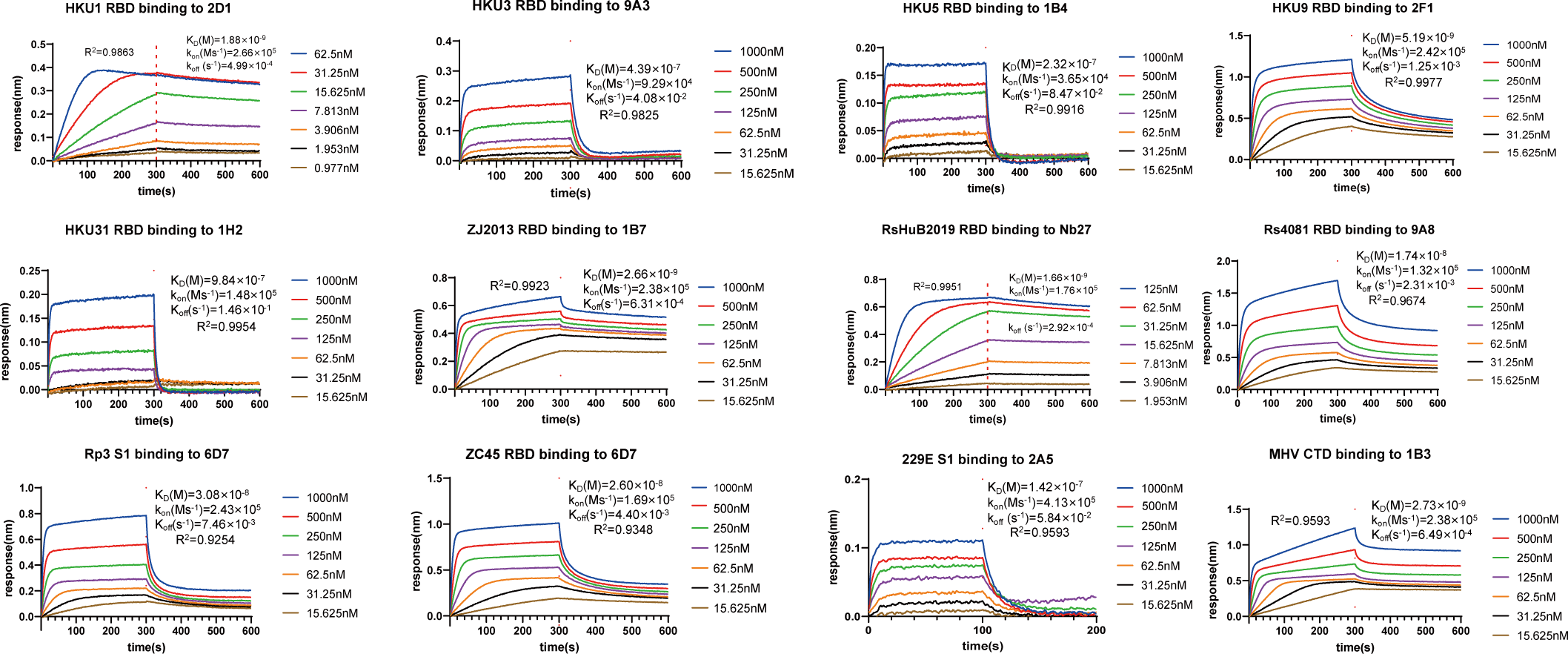
Binding kinetics between representative nanobodies and corresponding coronavirus antigens. Binding kinetics analyzed through BLI between representative nanobodies and the RBD or S1 of indicated coronaviruses.

**Extended Data Fig. 19.**
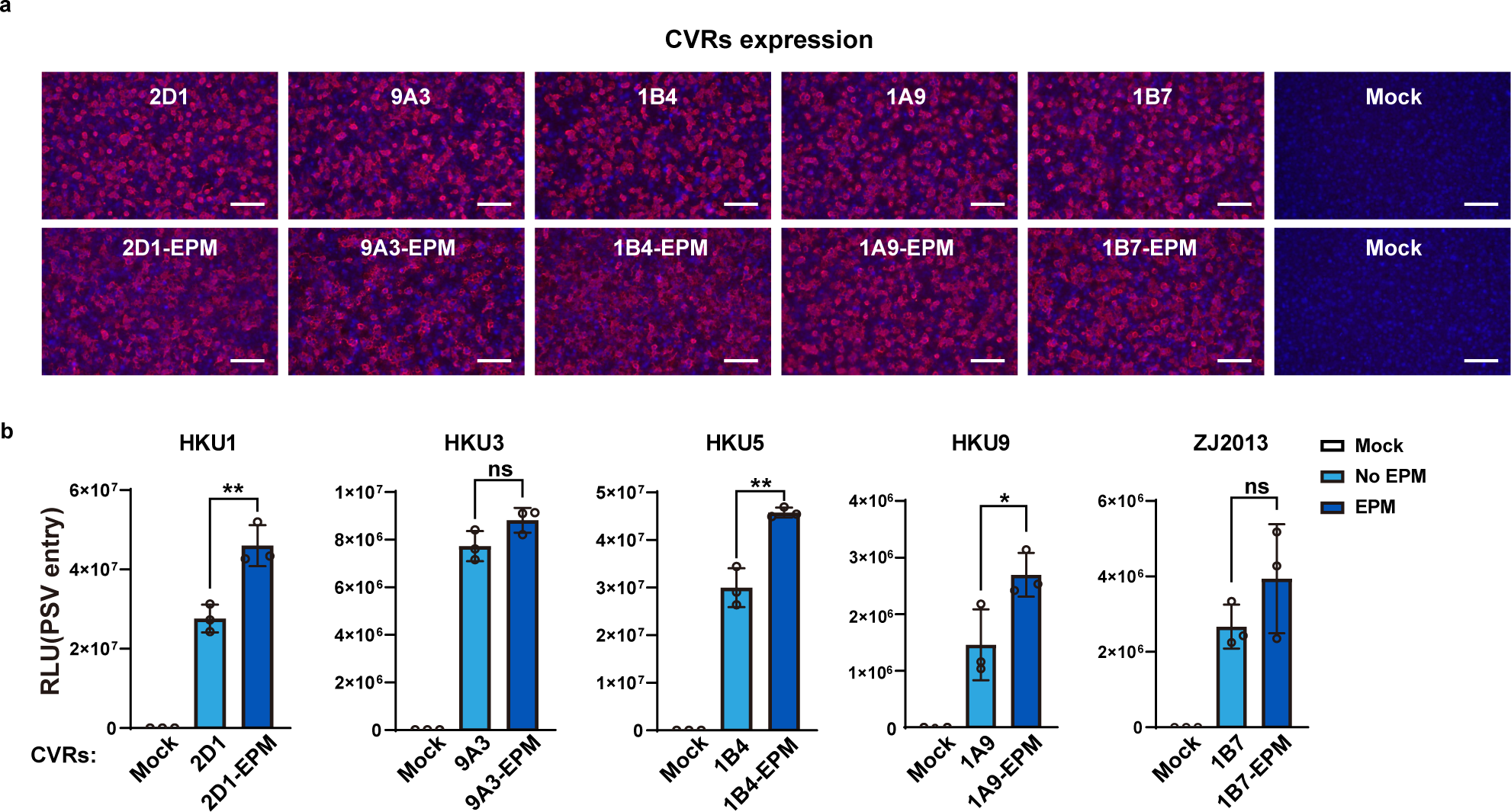
Comparison of the entry-supporting efficiency of several CVRs with or without the presence of EPM. **a,** Expression (**a**) and entry-supporting efficiency (**b**) of the indicated CVRs with or without EPM transiently expressed in the HEK293T cells. EPM: endocytosis prevention motif. Scare bars: 100μm. Unpaired two-tailed Student’s t-tests for **b**.

**Extended Data Fig. 20.**
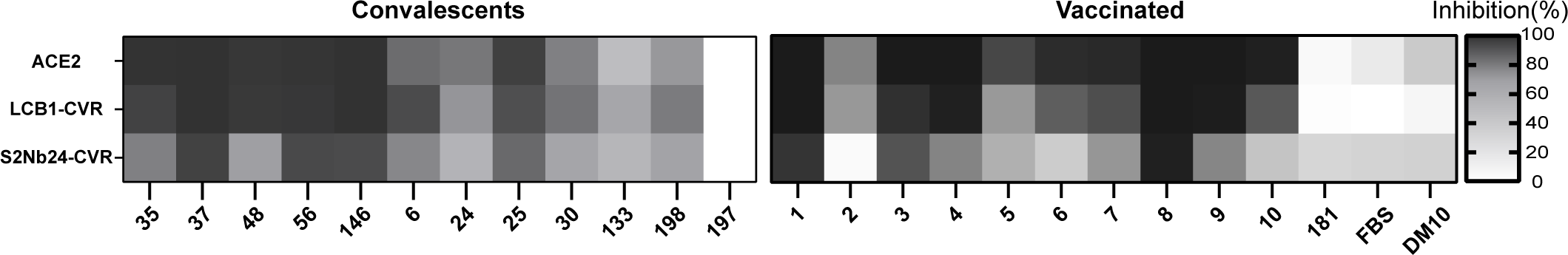
Comparison of sera neutralization activity using different infection models. Comparison of neutralization profiles of sera collected from COVID-19 convalescents (**a**) or vaccinated individuals (**b**) based on HEK293T cells expressing ACE2 or two different CVRs. Serum dilution: 1:200.

**Extended Data Fig. 21.**
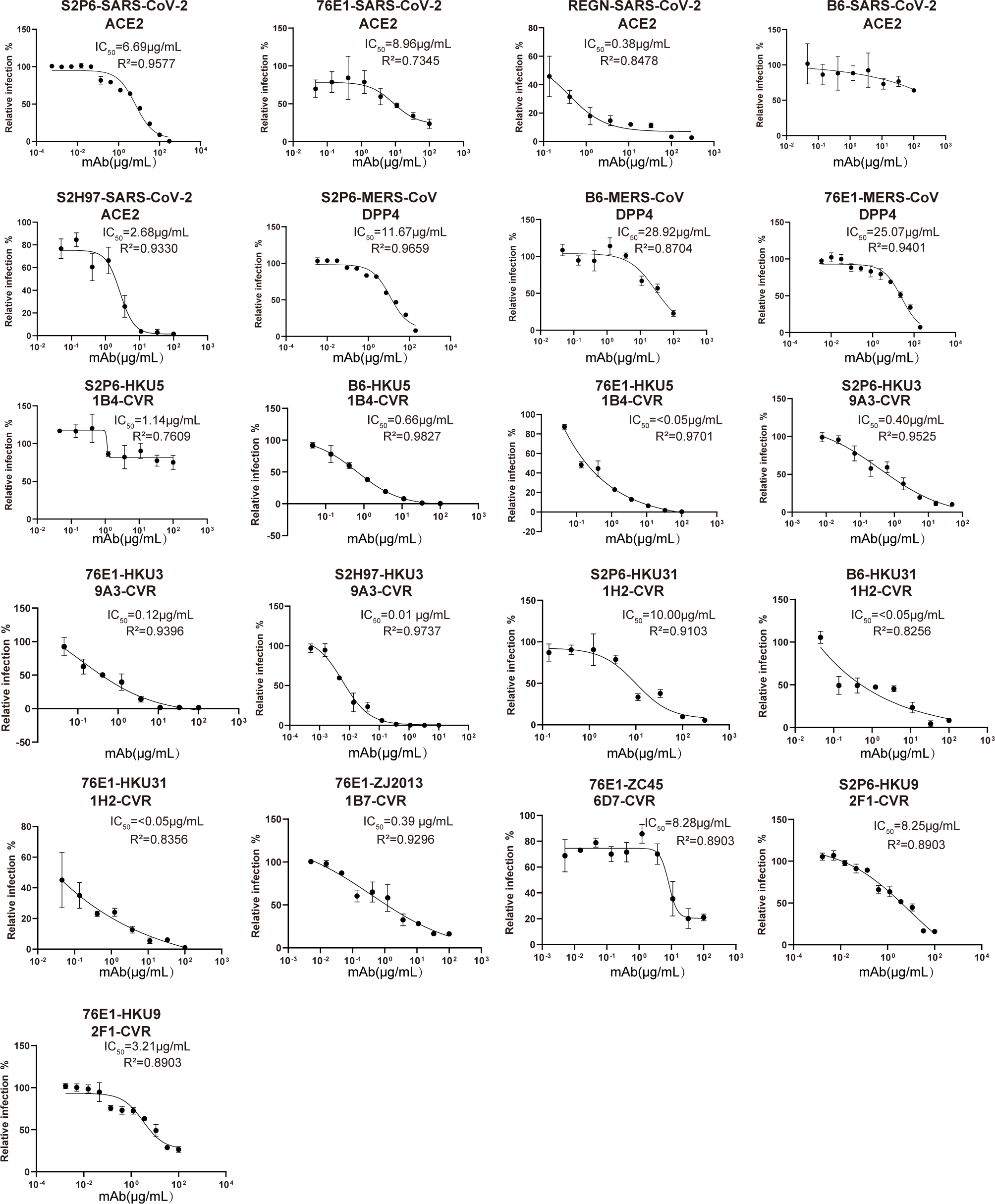
IC_50_ of selected broadly neutralizing antibodies against PSV entry of seven indicated coronaviruses supported by corresponding CVRs. Neutralization assays for each PSV were conducted in HEK293T stably expressing the indicated CVRs.

**Extended Data Fig. 22.**
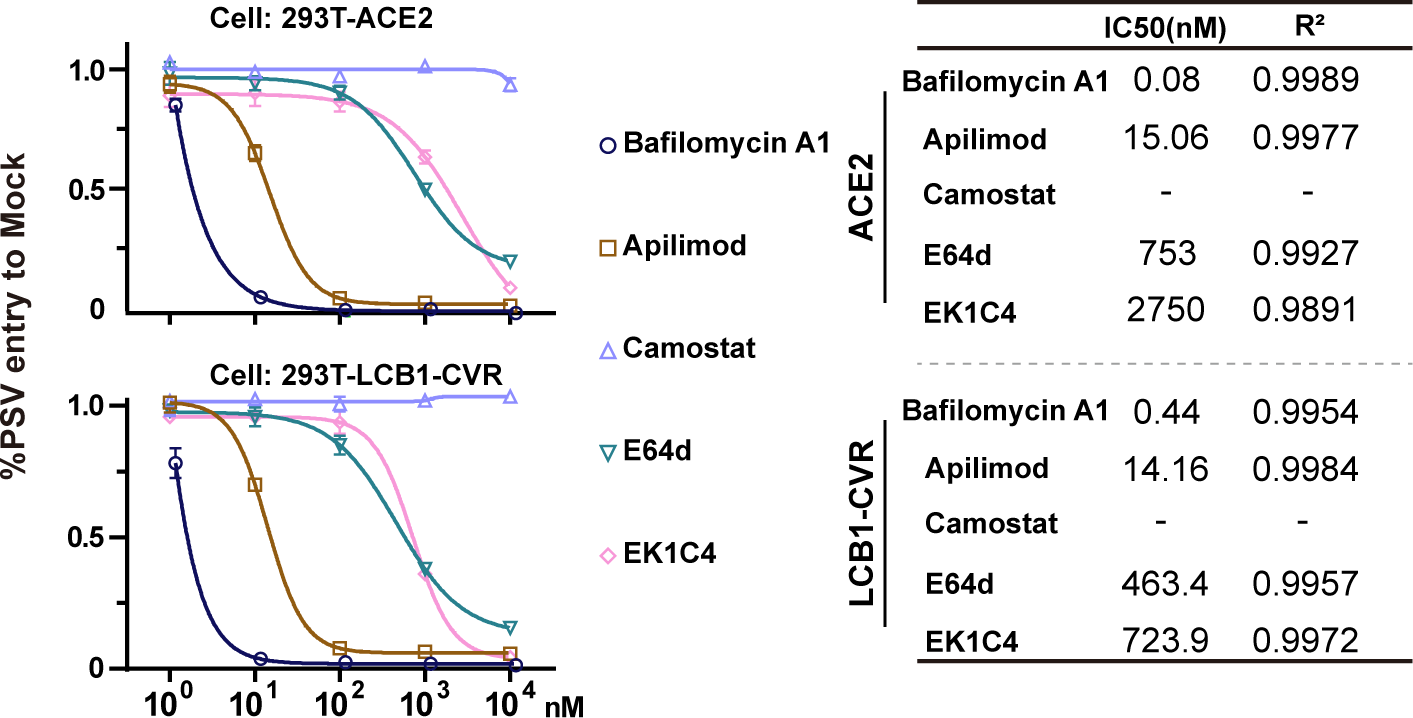
Inhibitory efficacy of entry inhibitors based on different infection models. The IC_50_ of selected entry inhibitors against SARS-CoV-2 PSV entry were determined in both HEK293T-ACE2 or HEK293T-LCB1-CVR cells.

**Extended Data Fig. 23.**
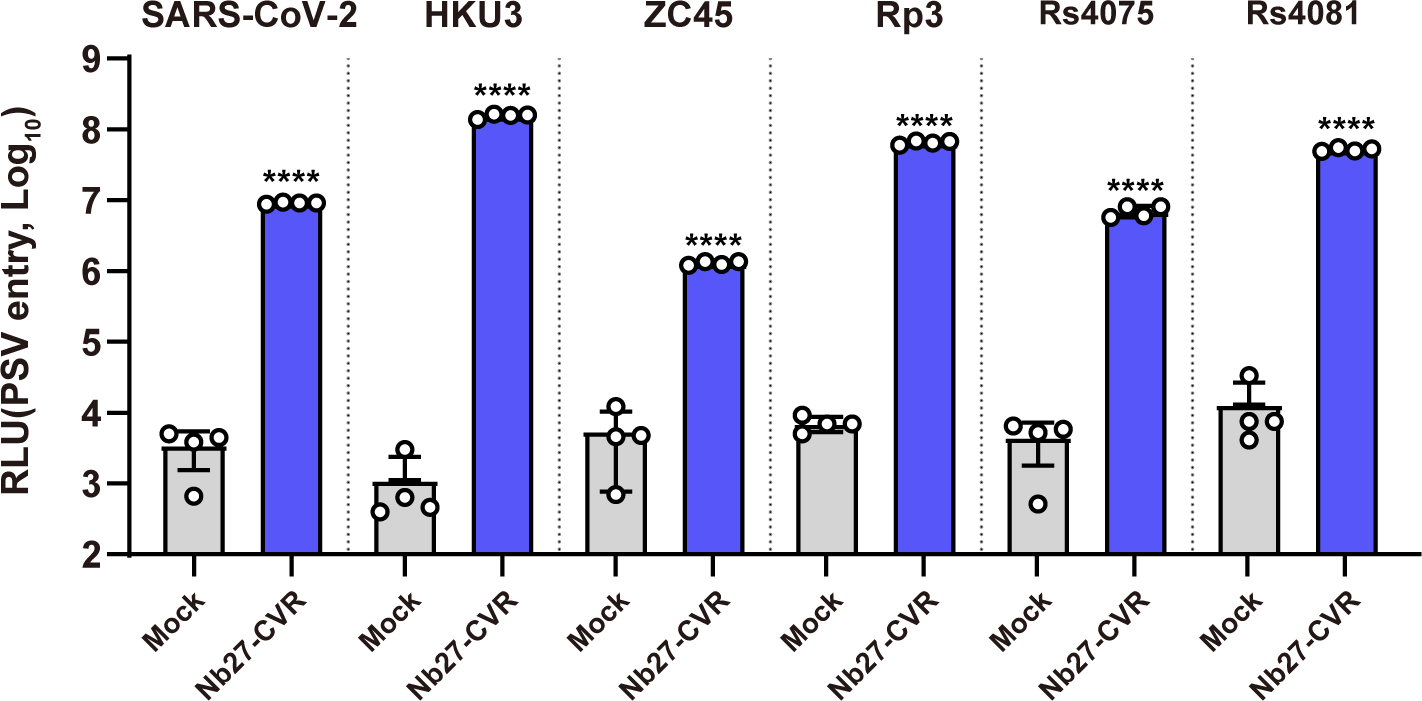
Pan-sarbecovirus entry-supporting ability of CVR-Nb27. The PSV entry-supporting ability of CVR-Nb27 was evaluated by six different sarbecoviruses in 293T cells. Unpaired two-tailed Student’s t-tests was employed for comparisons.

**Extended Data Fig. 24.**
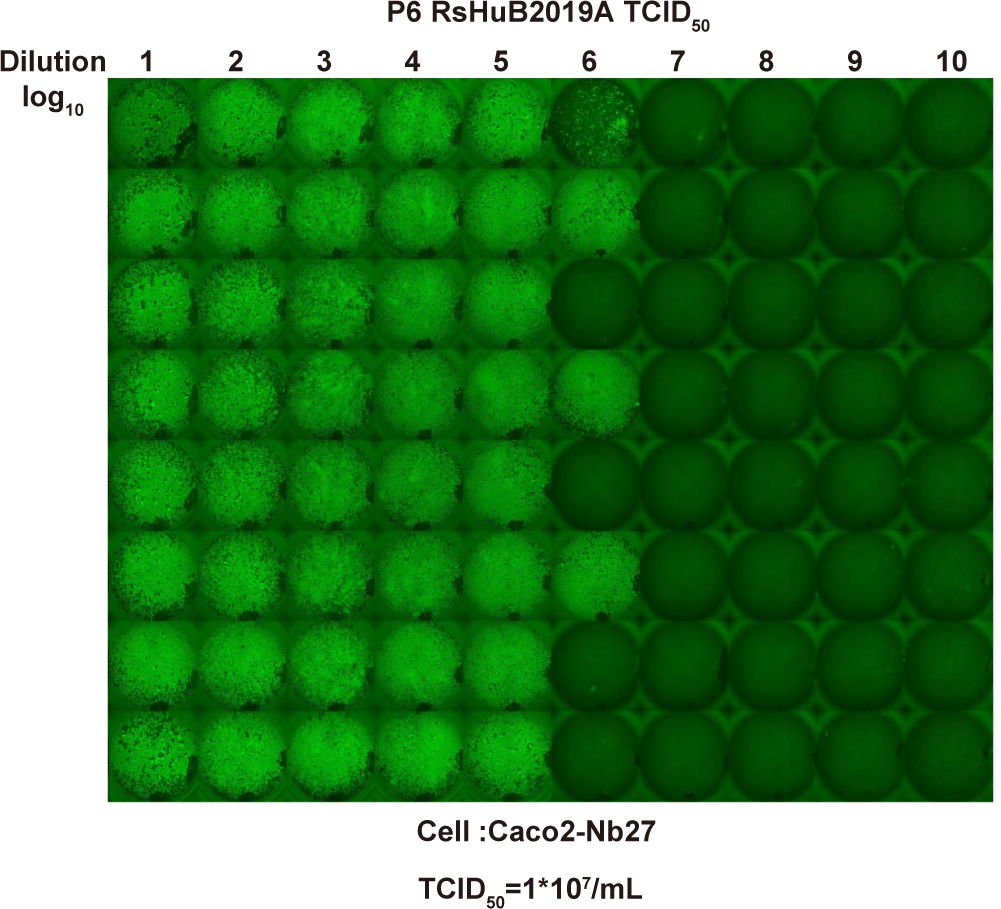
Demonstration of TCID_50_ determination assay for RsHuB2019A by Caco2-Nb27 cells. Caco2-Nb27 cells were inoculated with a 10-fold serial dilution of RsHuB2019A containing supernatant (Passage 6). The TCID_50_ was determined using immunofluorescence to detect the presence of N protein expression of the inoculated cells at 4 dpi, employing the Red-Muench method.

**Extended Data Fig. 25.**
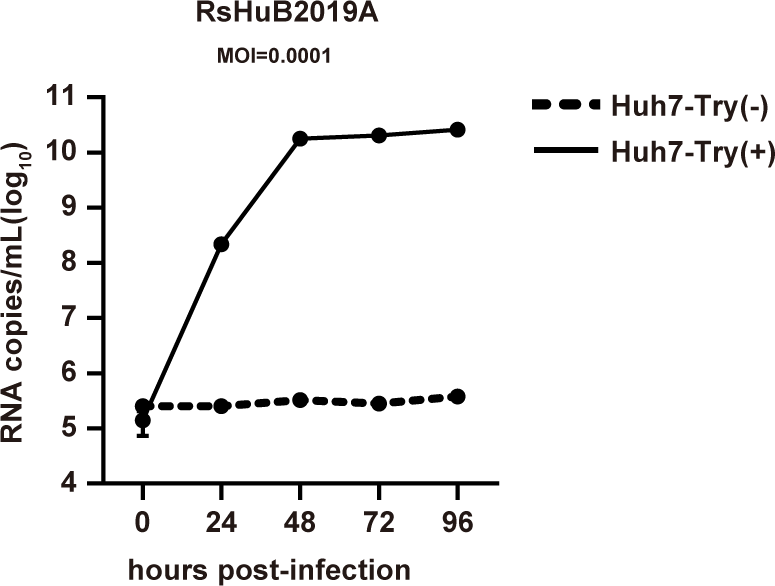
Trypsin-dependent propagation of RsHuB2019A in Huh-7 cells. The RsHuB2019A genomic RNA copies in the supernatant collected at indicated time points of infected Huh-7 cells were quantified by RT-qPCR using RdRP-specific primers. Inoculation was conducted at an MOI of 0.0001, with or without trypsin treatment. Try: Trypsin treatment.

**Extended Data Fig. 26.**
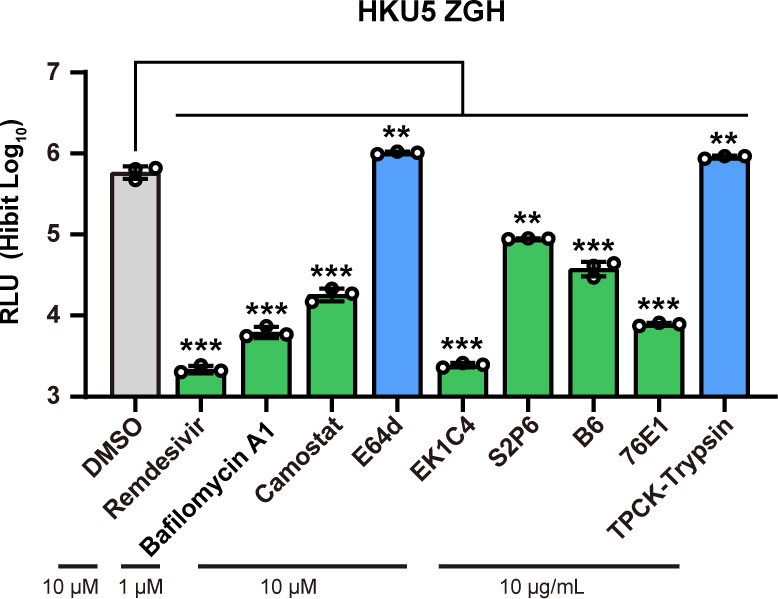
Inhibitory effect of selected anti-viral reagents against authentic HKU5-ZGH infection in Caco2-1B4. Inhibitors were coincubated with either the cells or the viruses for 1h and present in the culture medium during infection. The HiBit-based luciferase activity was determined at 48 hpi to assess the inhibitory effect of selected anti-viral reagents against the infection of authentic HKU5-ZGH in Caco2-1B4. Unpaired two-tailed Student’s t-tests was employed for comparisons.

**Extended Data Fig. 27.**
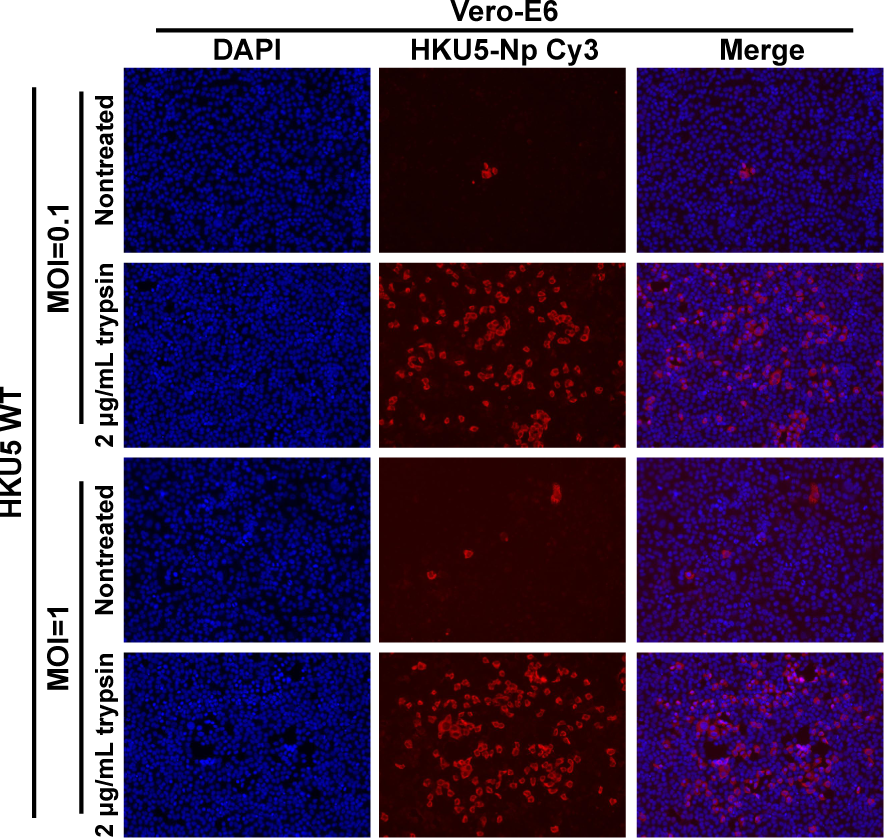
HKU5-WT propagation in Vero E6 cells with or without trypsin treatment. Vero E6 cells were infected with HKU5-WT at a multiplicity of infection (MOI) of 1 or 0.01, with or without the presence of trypsin at 2 μg/mL. The HKU5 infection efficiency was assessed using rabbit polyclonal antibodies targeting the HKU5 N protein(Cy3) at 48 hpi.

